# Glycine betaine-mediated transcriptional stimulation links its catabolism and vitamin B12 biosynthesis in *Rhizobiaceae*

**DOI:** 10.64898/2026.09.15.751800

**Authors:** Christian Rauch, Tamara Hoffmann, Marcel Wagner, Pornsri Charoenpanich, Jutta Gade, Fiona Ullmann, Konstantin Schneider, Hartwig Schröder, Oskar Zelder, Anke Becker

## Abstract

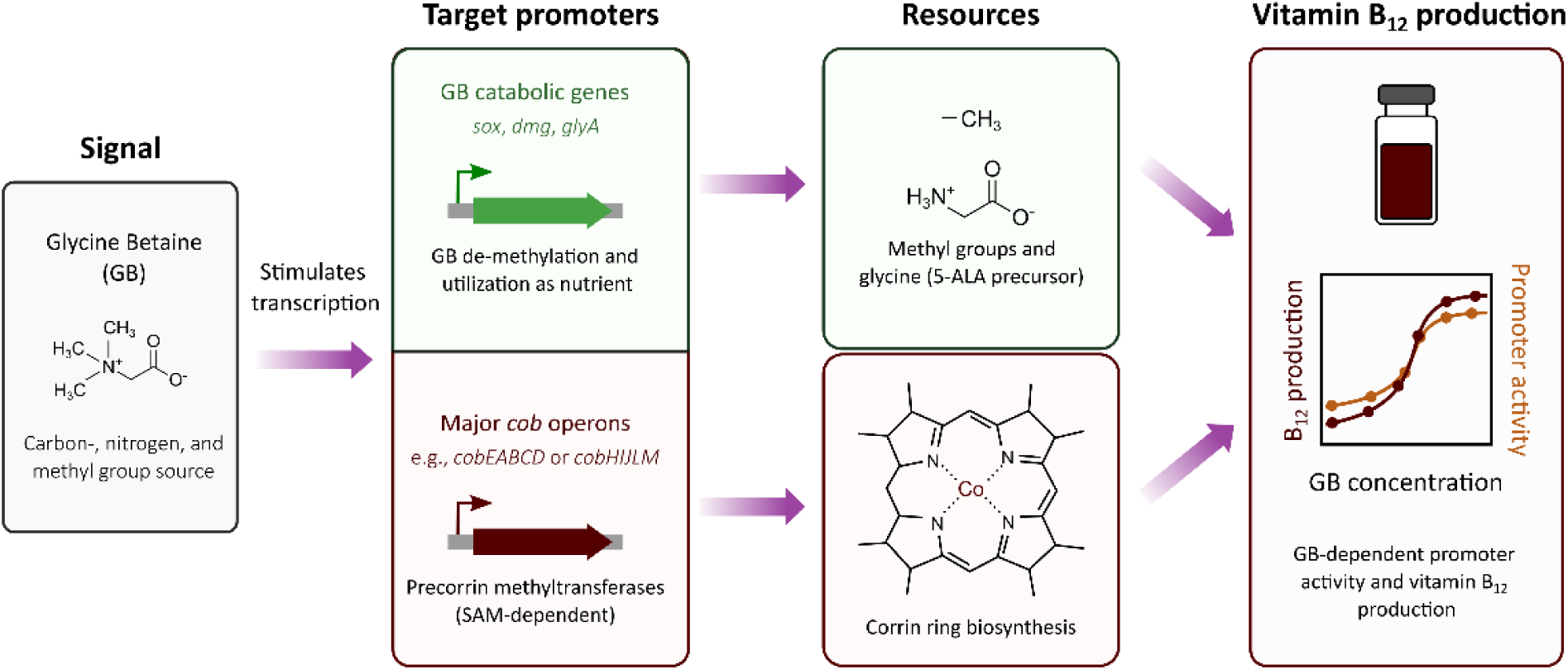

Glycine betaine (GB) is a trimethylated ammonium compound widely used as a compatible solute, and serves as a carbon, nitrogen, and methyl group source in many members of the *Rhizobiaceae*. GB is critical for high-level vitamin B₁₂ production, a cofactor synthesized exclusively by microorganisms. Vitamin B_12_ formation requires multiple methylation steps. In the industrial vitamin B_12_ production strain *Pseudomonas denitrificans*, GB has been shown to serve as a methyl group donor through its successive demethylation to glycine and to provide the carbon backbone for the universal porphyrin precursor 5-aminolevulinic acid. Moreover, previous proteomics studies have revealed increased levels of vitamin B₁₂ biosynthetic enzymes in the presence of GB. However, it has remained unclear at what level of gene expression this regulation occurs. Here, using promoter-reporter fusions in the vitamin B₁₂ overproducer *Ensifer adhaerens* LU2703 and in the low-producer *Sinorhizobium meliloti* 102F34, we demonstrate that GB activates promoters of both its own catabolic pathway and of major *cob* operons. Two of these operons encode the complete set of *S*-adenosylmethionine-dependent methyltransferases required for corrin ring biosynthesis, the foundational, rate-limiting stage in vitamin B_12_ production. Stimulation of *cobE* and *cobH* promoters as well as vitamin B₁₂ production yields followed sigmoidal dose-response curves with saturation in a similar concentration range, suggesting a tight coupling between GB supply and biosynthetic output. Our findings uncover a novel regulatory layer in vitamin B₁₂ biosynthesis and provide promoter-probe set-ups for further exploration of GB-mediated transcriptional control.

## Introduction

Glycine betaine (GB) is a trimethylated organic osmolyte that is ubiquitously found in organisms across all kingdoms of life. In microorganisms as well as animals and plants it is used as a powerful protectant against osmotic stress and growth temperature extremes and, via its degradation, as an energy, nitrogen and carbon source (Boncompagni *et al*., 1999; Diamant *et al*., 2003; Barra *et al*., 2006; Burg and Ferraris, 2008; Zou *et al*., 2016; Bremer and Krämer, 2019). It is not only relevant for protecting cells in natural settings but has also emerged as an interesting substance for biotechnological applications. As a chemical chaperone, it stabilizes macromolecules like membranes and proteins both *in vitro* and *in vivo* (Burg and Ferraris, 2008). For example, supplementation of industrial fermentation media with GB has been shown to enhance cell growth in high-osmolarity media, to improve the solubility and stability of proteins and enzymes, and to serve as a methyl donor in bioproduction processes (reviewed by Zou *et al*., 2016).

In response to elevated external osmolarity, many organisms scavenge GB from the environment via specialized high-affinity transport systems. They accumulate it up to molar concentrations in order to balance the osmotic gradient between their cytoplasm and the environment (Wood *et al*., 2001; Bremer and Krämer, 2019). If preformed GB is not available, several microorganisms and many plants can synthesize GB from its precursor choline via a two-step oxidation pathway. Notably, the capability to import or synthesize GB from its precursor does not necessarily lead to an intracellular accumulation of the compound since in some bacteria: particularly within the *Rhizobiaceae*, choline oxidation to GB represents the initial step for its subsequent catabolism rather than its retention as an osmoprotectant (Smith *et al*., 1988; Talibart *et al*., 1997). Many salt-tolerant rhizobial strains accumulate GB as osmoprotectant only transiently in the early exponential growth phase before it is degraded and used as a nutrient (Bernard *et al*., 1986; Boncompagni *et al*., 1999).

In the rhizobial model species *Sinorhizobium meliloti,* GB uptake is mediated by three transport systems: the high affinity BCCT importer BetS, the low affinity ABC transporter Hut, and the Opp-type transporter Prb, all of which are active under iso-osmotic conditions and serve to import GB as a source of carbon and nitrogen (Bernard *et al*., 1986; Smith *et al*., 1988; Boscari *et al*., 2002). Of these, only BetS is additionally activated in response to hyperosmotic stress (Boscari *et al*., 2002). Uptake of the precursor choline is mediated by the highly specific ABC transporter Cho, which is substrate-inducible and not further induced by osmotic stress (Dupont *et al*., 2004).

Once choline is taken up by *S. meliloti*, it is oxidized to GB. The responsible genes are encoded by the *betICBA* operon which comprises both the gene for the choline-responsive repressor, *betI*, and the three structural genes, *betC*-*betB*-*betA*, that catalyze the choline to GB conversion. It has been shown that BetI represses *betICBA* transcription via binding to the BetI box located in the *betI* promoter region (Mandon *et al*., 2003). In the presence of choline in the growth medium the promoter region is liberated from its repressor and transcription takes place. Expression of the GB synthesis pathway is thereby substrate-induced but not osmotically regulated (Mandon *et al*., 2003).

Via its catabolism, GB serves as nitrogen and carbon source and can provide methyl groups for biosynthetic pathways in *S. meliloti* and other *Rhizobiaceae* (Boncompagni *et al*., 1999; Barra *et al*., 2006). Its degradation starts with sequential demethylation of its fully methylated nitrogen head group to form glycine. The intermediates are dimethyl glycine (DMG) and sarcosine (monomethyl glycine) (Fig. 1A). At least the first demethylation step is linked to vitamin B_12_ synthesis. This reaction is catalyzed by betaine-homocysteine *S*-methyltransferase (BHMT) which transfers the methyl group to homocysteine to form methionine, thereby fueling the *S*-adenosyl methionine (SAM)-cycle that provides methyl groups for SAM-dependent methylation reactions (White and Demain, 1971; Barra *et al*., 2006). After full demethylation, the resulting glycine can be converted into serine via GlyA. Deamination of serine catalyzed by L-serine dehydratase (SDA) would subsequently produce ammonia and pyruvic acid that will be introduced into the central metabolism. Alternatively, glycine can be used by 5-aminolevulinic acid (5-ALA) synthase (HemA) to form 5-ALA, the essential precursor for tetrapyrrole biosynthesis, and therefore for vitamin B_12_ production in *S. meliloti*.

**Fig. 1:**
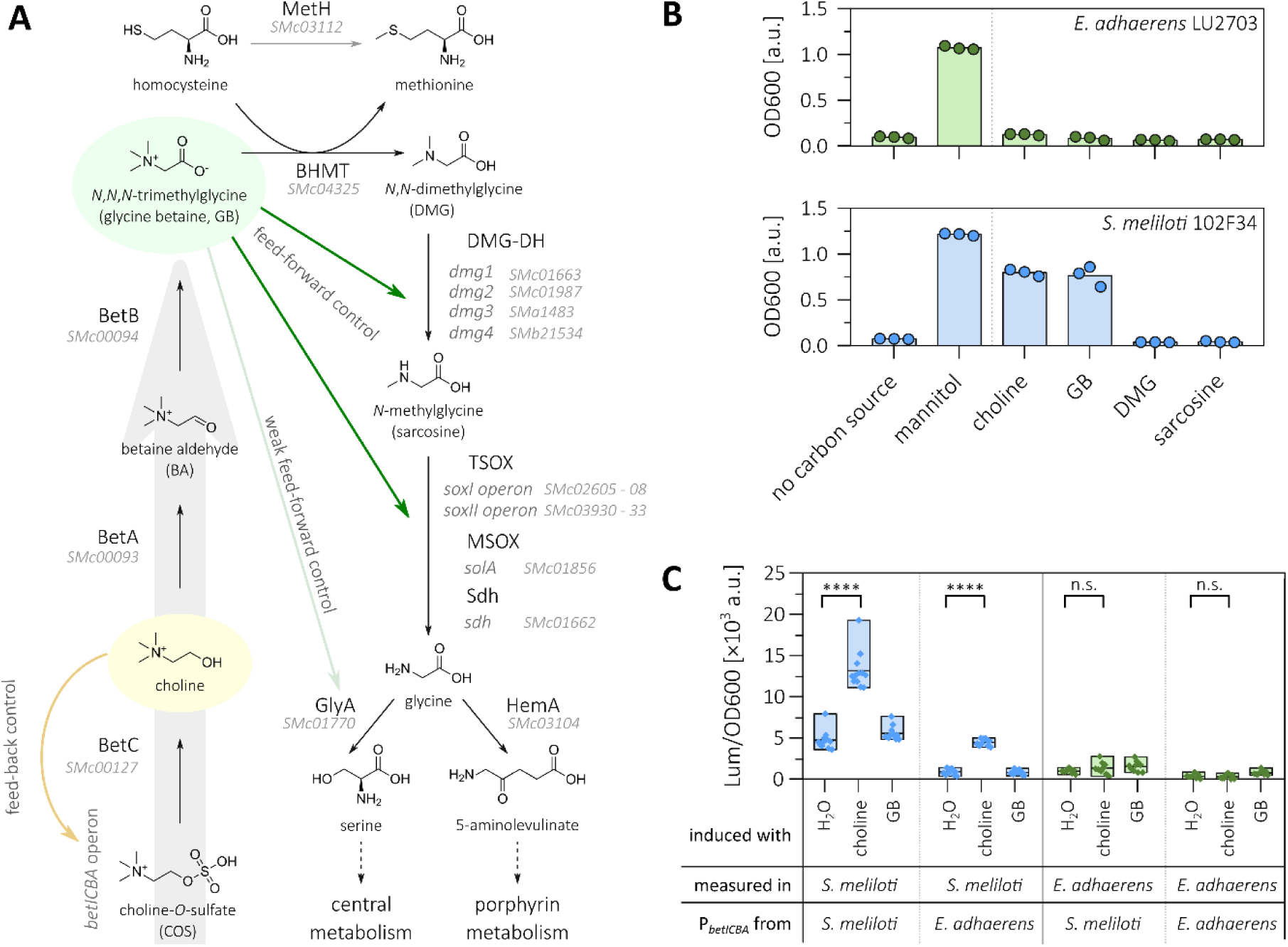
Metabolism, nutrient utilization, and transcriptional impact of GB in *E. adhaerens* LU2703 and *S. meliloti*. **(A)** Synthesis and degradation pathway of GB in *S. meliloti* 102F34. The *betICBA* operon, encoding proteins that are highlighted by the grey arrow, is transcriptionally activated by choline (Mandon *et al*., 2003) (yellow arrow). GB exerts a feed-forward control on its own degradation pathway by affecting specific genes downstream in the pathway (green arrows). **(B)** Growth of *S. meliloti* 102F34 (bottom, blue bars and circles) and *E. adhaerens* LU2703 (top, green bars and circles) after 48 h in MOPS-buffered minimal medium using either mannitol, GB, or metabolic precursors and products of GB as sole carbon source. Cultures were grown in microplates and measured on a Tecan Infinite M Plex reader. Precultures were grown in TY medium supplemented with 600 µg/mL streptomycin and washed twice with minimal medium without carbon source before inoculation. **(C)** Transcriptional-translational promoter-probe assay characterizing the promoter of the *betICBA* operon from *E. adhaerens* LU2703 and *S. meliloti* 102F34 (399 bp and 290 bp upstream of the respective *betI* start codon, respectively) grown in MOPS-buffered minimal medium in the presence of either 1 mM choline, 1 mM GB, or H_2_O (no inducer) in microplates and measured on a Tecan Infinite M200 Pro reader. Data shown 6 h after addition of the putative inducers. Data are based on 8 to 12 biological replicates. Statistical significance was calculated by multiple *t* test (n.s.: not significant, ****: *p_adj_* < 0.000 01).

In *S. meliloti*, the GB catabolic route is active under iso-osmotic conditions and is further stimulated by GB or its precursor choline, indicating a classical feed-forward regulation (Smith *et al*., 1988). Conversely, osmotic stress strongly suppresses catabolic enzyme activities even in the presence of GB or choline, thereby enabling osmoprotective GB accumulation, at least transiently during early exponential growth (Bernard *et al*., 1986; Smith *et al*., 1988).

Cobalamin (Vitamin B_12_) is one of nature’s most complex non-polymeric molecules, essential for humans and other animals but synthesized exclusively by certain bacteria and archaea. The cobalamin molecule is characterized by a highly methylated porphyrin ring, coordinating a central cobalt ion, a cyano-(vitamin B_12_), methyl-, hydroxy- or adenosyl group as upper ligand and a conserved lower axial ligand, 5,6-dimethylbenzimidazole (DMBI). For simplicity, we refer to cobalamins collectively as vitamin B_12_. The aerobic vitamin B_12_ biosynthetic pathway, best characterized in *Pseudomonas denitrificans* (nom. rej.), comprises approximately 25 to 30 enzymatic steps divided into three major stages: (i) assembly and methylation of the corrin ring from uroporphyrinogen III, (ii) cobalt insertion and amidation of the corrinoid intermediate, and (iii) nucleotide loop assembly including synthesis and attachment of the lower ligand (reviewed by Martens *et al*., 2002; and Balabanova *et al*., 2021). The corresponding *cob* genes are typically organized in strain-specific chromosomal clusters related to these stages. Transcription of *cob* genes often takes place in operons, thereby enabling their coordinated transcriptional control, often involving cobalamin-responsive riboswitches. Although gene content and cluster arrangement vary among producer strains, the biosynthetic enzymes are well conserved, and recombinant genes from a small number of characterized strains, including those of *S. meliloti*, have been used to construct new recombinant vitamin B_12_ overproducer strains (Martens *et al*., 2002; Balabanova *et al*., 2021).

Early studies on vitamin B_12_ fermentation in the overproducer *P. denitrificans* showed that GB was necessary for high vitamin B_12_ production rates (Demain *et al*., 1968). Multiple studies support this, and the prevailing explanation for the beneficial effect of GB in vitamin B_12_ production is its function as a source of methyl groups and carbon backbone for the essential tetrapyrrol precursor, 5-ALA (White and Demain, 1971; Martens *et al*., 2002; Li *et al*., 2008; Xia *et al*., 2015). However, pioneering studies already hinted at a potential regulatory role of GB during vitamin B_12_ production (Demain *et al*., 1968; White and Demain, 1971) due to the finding that GB was essential for vitamin B_12_ production and could not be replaced by other methyl group donors in *P. denitrificans*. While early investigations showed a relationship between the presence of GB and the specific activity of HemA (Kusel *et al*., 1984), more recent data suggest a broader change in protein abundances of enzymes involved in the vitamin B_12_ synthetic network in response to GB in the medium (Li *et al*., 2022).

Here, we report that the novel vitamin B_12_ overproducer strain, the Alphaproteobacterium *Ensifer adhaerens* LU2703, responds to GB by transcriptionally activating its degradation pathway genes and genes involved in vitamin B_12_ biosynthesis. Similar regulatory patterns observed in the related wild type strain *S. meliloti* 102F34, a vitamin B_12_ low-producer, suggest that this mechanism may be conserved among major producer strains, including *P. denitrificans*. Our discovery of the GB-dependent transcription level regulatory link between vitamin B_12_ biosynthesis and GB metabolism deepens our understanding of the regulatory principles governing the production of this important vitamin, thereby providing new starting points for the genetic optimization of production strains.

## Results

### *E. adhaerens* LU2703 contains genes related to GB uptake, synthesis and catabolism but does not use GB as carbon source for growth

It has been shown that members of the order Hyphomicrobiales (formerly Rhizobiales) are natural overproducers of vitamin B_12_, particularly strains belonging to the species *E. adhaerens* (Vu *et al*., 2013; Zhao *et al*., 2019; Bampidis *et al*., 2023; Liu *et al*., 2023)and *S. meliloti* (Dong *et al*., 2016). To investigate GB-dependent regulation of GB metabolism and vitamin B_12_ biosynthesis, we focused on the novel, patent-restricted (Becker *et al*., 2026b, 2026a) vitamin B_12_ overproducer *E. adhaerens* LU2703. As a first step, we sequenced and annotated the genome of *E. adhaerens* LU2703. It consists of three replicons, a chromosome of 3 951 890 bp and two large extrachromosomal replicons of 1 784 594 bp and 1 592 480 bp. This genomic architecture is highly reminiscent of that of *S. meliloti* (Galibert *et al*., 2001) which also has a tripartite genome consisting of a main chromosome, a chromide and a megaplasmid. The 7.3 Mbp *E. adhaerens* LU2703 genome contains 6854 putative protein-coding genes and shares an average nucleotide identity (Yoon *et al*., 2017; Chalita *et al*., 2024) of 99.11 % with the genome sequence of the type strain *E. adhaerens* Casida A (Williams *et al*., 2017) and 79.29 % with *S. meliloti* 102F34 (Jozefkowicz *et al*., 2017), a vitamin B_12_ low-producer ((5.06 ± 0.05) mg B12/L after 5 days compared to (38 ± 2) mg B12/L produced by *E. adhaerens* LU2703, Supplementary Data S1).

In order to characterize the genetic capabilities of *E. adhaerens* LU2703 to import, synthesize, and degrade GB, we annotated the corresponding genes based on *S. meliloti* 102F34 genes encoding proteins that have been shown to be involved in GB metabolism (all genes, gene products and their identities are summarized in Table S1). For GB and choline uptake, *E. adhaerens* LU2703 encodes putative transport systems including the choline-specific ABC transporter Cho (Dupont *et al*., 2004), the Opp-type transporter Prb known to transport proline betaine as well as choline or GB (Alloing *et al*., 2006), and the low affinity GB importer Hut (Boncompagni *et al*., 2000). Notably, we could not identify a homolog of BetS, which is the only osmotically induced GB transporter in *S. meliloti* (Boscari *et al*., 2002). A putative *betICBA*-operon, which is widespread within Alphaproteobacteria, encodes the putative GB synthesis pathway starting from the precursors COS (choline-*O*-sulfate) or choline. For catabolism we identified genes for the full demethylation-cascade: (i) a putative betaine homocysteine *S*-methyltransferase (BHMT; 88.4 % sequence identity to the corresponding protein in *S. meliloti* 102F34) coupling the first de-methylation reaction to methionine synthesis thereby producing DMG, (ii) four DMG dehydrogenase (DMG-DH) homologs (85.5 to 94.4 % and 43.6 % sequence identity to Dmg1, Dmg2, Dmg4 and Dmg3, respectively) (Table S1) mediating conversion of DMG to sarcosine, and (iii) the putative tetrameric sarcosine oxidases (TSOX) SoxI (40.1 % to 76.2 % identity) and SoxII (79.9 % to 98.1 % identity), the putative monomeric sarcosine oxidase (MSOX) SolA (73.5 % identity), and the putative sarcosine dehydrogenase Sdh (91.4 % identity) producing glycine (Table S1). Gene products associated with glycine utilization are highly conserved between both species. These enzymes catalyze reactions channeling glycine either into porphyrin biosynthesis via 5-ALA synthase (HemA) activity, or into central metabolism via serine hydroxymethyltransferase GlyA activity and the glycine cleavage system GcvTPH.

Since our *in silico* analysis suggested that *E. adhaerens* LU2703 possesses a GB catabolic route, we tested its ability to utilize GB or GB-related compounds as sole carbon source. *S. meliloti* 102F34, known for its ability to catabolize GB, was used as a control (Smith *et al*., 1988). The strains were cultivated for 48 h in minimal medium containing GB or related compounds as sole carbon sources, each normalized to equivalent molar carbon concentrations. Both strains could not grow in carbon-free minimal medium but grew efficiently on mannitol, which served as a positive control (optical density at 600 nm (OD600) of 1.0 to 1.2 after 48 h) (Fig. 1B). Unlike *S. meliloti* 102F34, *E. adhaerens* LU2703 failed to grow with GB or choline as sole carbon source (Fig. 1B). Neither strain grew in minimal medium supplemented with DMG or sarcosine as sole carbon source. Repeating the experiment in the presence of mild osmotic stress (100 mM NaCl), to account for possible osmotic induction of GB uptake systems, did not alter the growth pattern (Fig. S1).

### GB stimulates promoter activities of GB metabolism-related genes in *E. adhaerens* LU2703 and *S. meliloti* 102F34

To assess whether GB functions as an effector molecule in the transcriptional regulation of the GB metabolic network, we identified the promoters of this network and analyzed their activities in response to GB. A Cappable-seq analysis (Ettwiller *et al*., 2016) in *E. adhaerens* LU2703 was performed to determine transcriptional start sites (TSS). We mapped a total of 9948 TSS (Table S2) above a defined quality threshold (for details see Experimental procedures), allowing identification of promoters for all genes and operons of the GB metabolic network (Fig. S2-S5).

In *Rhizobiaceae*, the *bet* genes encoding enzymes for GB biosynthesis are organized within the *betICBA* operon, whose transcription is controlled by the choline-responsive repressor BetI (Mandon *et al*., 2003). The promoter and 5’-UTR of this operon, the BetI binding site and BetI itself show only slight differences between *E. adhaerens* LU2703 and *S. meliloti* 102F34 (Fig. S6, Table S1). To assess whether the *betI* promoter in both strains share the same regulation pattern, we fused the *betICBA* upstream regions (399 bp and 290 bp, respectively) with the *luxCDABE* reporter operon on a single-copy plasmid (Meier *et al*., 2024). The resulting transcriptional-translational fusion constructs were then introduced both into *E. adhaerens* LU2703 and *S. meliloti* 102F34, allowing promoter activities to be assessed in *E. adhaerens* LU2703 and *S. meliloti* 102F34 for each construct (Fig. 1C). The resulting reporter strains were grown in minimal medium with either 1 mM choline or 1 mM GB, or no supplement. After 6 h of incubation in *S. meliloti* 102F34, the native P*_betI-Sm_* promoter showed higher basal reporter activity ((4.7 ± 1.1) × 10^3^ Lum/OD600) than the heterologous P*_betI-Ea_* promoter ((0.8 ± 0.3) × 10^3^ Lum/OD600) in the absence of either GB or choline. Adding GB did not influence these basal levels. As expected, activities of both promoters increased in *S. meliloti* when choline was added to the medium (5.1-fold for P*_betI-Sm_* and 2.8-fold for P*_betI-Ea_*), indicating de-repression through formation of a BetI-choline complex. However, when *E. adhaerens* LU2703 was the host, both promoter-reporter fusions exhibited only low basal activities (P*_betI-Sm_* (0.8 ± 0.2) × 10^3^ Lum/OD600 and P*_betI-Ea_* (0.3 ± 0.3) × 10^3^ Lum/OD600) and addition of either choline or GB to the medium did not significantly increase these activities. This was also the case for the promoter-reporter fusions in an *E. adhaerens* LU2703 *betI* deletion mutant complemented with the *S. meliloti* 102F34 *betICBA* operon on a single copy plasmid, while the *betI* deletion mutant carrying the empty plasmid showed increased P*_betI_* reporter activities (Fig. S7). The most likely explanation for this difference would be insufficient import of choline by *E. adhaerens* LU2703.

In *S. meliloti*, it has been shown that the fate of synthesized or imported GB is its degradation (Talibart *et al*., 1997). We therefore analyzed the response of promoters of GB catabolic genes to GB. To this end, the corresponding *E. adhaerens* LU2703 and *S. meliloti* 102F34 promoter regions (approximately 300 to 400 bp upstream of the start codons) were fused to a *luxCDABE* reporter cassette on a single copy plasmid (Table S3), and introduced into their cognate wild type strain. These reporter strains were grown in minimal medium with and without 1 mM GB. The response of the promoters to the presence of GB in both *E. adhaerens* and *S. meliloti* are shown in Fig. 2. Promoters of genes and operons putatively involved in GB de-methylation reactions, namely *dmg2* and the *soxII* operon (Fig. 1A), showed high GB-mediated stimulation. Both Dmg2 and the SoxII complex showed the highest degree of sequence similarity for their respective enzymatic functions (Table S1) between both species reflecting the shared regulatory pattern. In comparison, the *glyA* promoters also displayed enhanced activities in response to GB, although their average increase was lower in both *E. adhaerens* LU2703 and *S. meliloti* 102F34. This may be due to the high basal activities measured in the *glyA* promoter-reporter strains: in *E. adhaerens* LU2703, P*_glyA_* activity rose from (8 ± 2) × 10^3^ Lum/OD600 to (20 ± 7) × 10^3^ Lum/OD600 and in *S. meliloti* 102F34 from (29.2 ± 1.8) × 10^3^ Lum/OD600 to (59 ± 7) × 10^3^ Lum/OD600. In both species, this marks the highest values of reporter activities for promoters related to the GB metabolic network in both the induced and non-induced states.

**Fig. 2:**
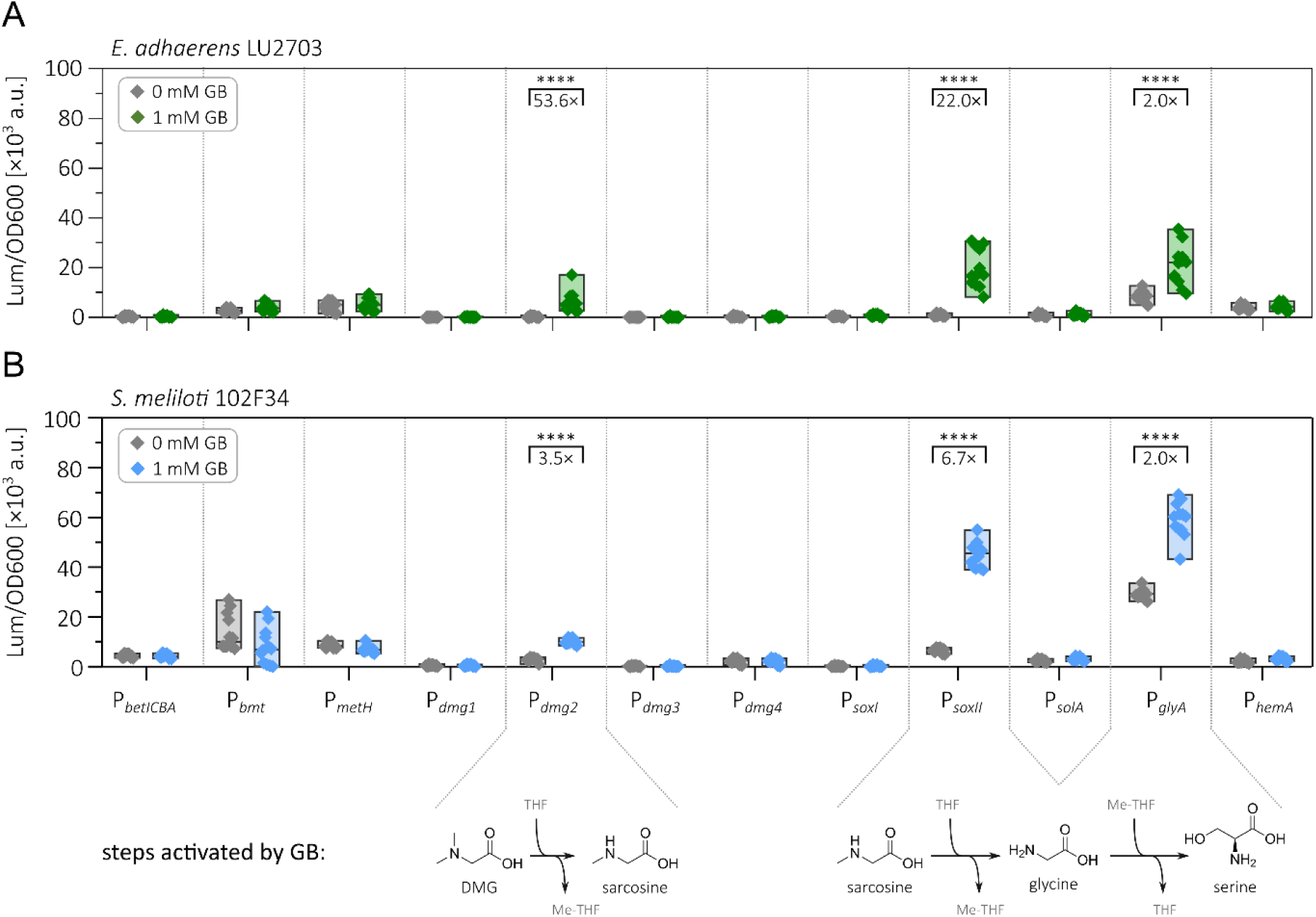
Activities of promoters involved in GB metabolism and their response to exogenous GB. Transcriptional-translational promoter-probe assays of promoters of genes involved in the GB metabolic network of *E. adhaerens* LU2703 **(A)** and *S. meliloti* 102F34 **(B)**. Strains carrying transcriptional-translational promoter-*luxCDABE* reporter constructs on a single-copy number plasmid were cultured in MOPS-buffered minimal medium with 0 mM or 1 mM GB for 6 h in microplates before data was taken with a Tecan Infinite M200 Pro reader. Data shown is based on 6 biological replicates each measured twice on different days. Statistical significance was calculated by multiple *t* test (****: *p_adj_* < 0.000 01, non-significant results were not labeled).

### *E. adhaerens* LU2703 shows GB-dependent vitamin B_12_ production and harbors a *cob* gene repertoire similar to *S. meliloti* 102F34

High-level vitamin B_12_ production requires GB availability and its degradation, as both methyl groups for corrin ring methylation and precursors for the tetrapyrrole core biosynthesis are derived from GB. Since GB up-regulates the transcription of the genes involved in its own catabolism in the vitamin B_12_ overproducer *E. adhaerens* LU2703, we hypothesized that GB may also influence the transcriptional regulation of vitamin B_12_ synthesis genes, thereby enhancing vitamin B_12_ production.

To assess the dependence of vitamin B_12_ production on GB, *E. adhaerens* LU2703 was cultivated in shaking flasks in MOPS-buffered fermentation medium (Table S4) supplemented with increasing concentrations of GB. After five days of incubation, when cultures reached OD600 values between 20 and 32.5, vitamin B_12_ concentrations were quantified by HPLC (Fig. 3). Cultures grown without GB contained only low amounts of vitamin B_12_, whereas supplementation with GB resulted in maximal yields of 39 mg/L, corresponding to a 116-fold increase. Vitamin B₁₂ production followed a sigmoidal dose-response relationship with increasing GB concentrations and reached a plateau at GB concentrations above 51 mM (Fig. 3). These results suggest that GB availability is rate-limiting for vitamin B_12_ synthesis at low concentrations, whereas other factors become limiting under GB-saturated conditions. The observed GB-dependence is consistent with GB acting as methyl group and precursor donor but also with an additional role as regulatory signal. As a basis for investigating the latter possibility, we analyzed the genetic make-up of *E. adhaerens* LU2703 related to vitamin B_12_ biosynthesis.

**Fig. 3:**
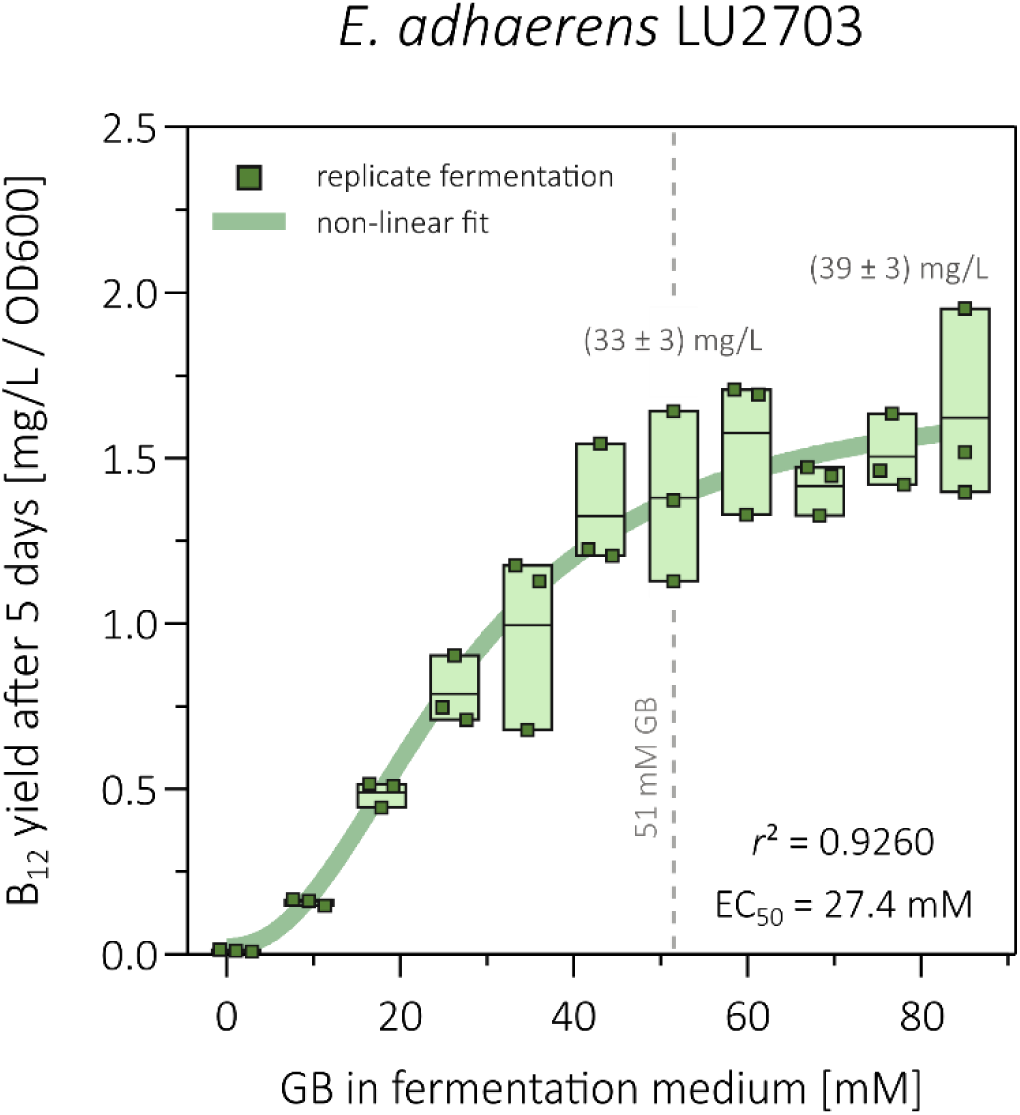
GB concentration-dependent production of vitamin B_12_ by *E. adhaerens* LU2703. The strain was grown in MOPS-B_12_ medium supplemented with 0 to 80 mM GB under microoxic conditions in narrow-necked shaking flasks for five days. Green rectangles show vitamin B_12_ yield of cultures normalized to OD600 (measurement details in Experimental procedures). The line within boxes represents the average yield based on three replicate batch fermentations. The green line represents a non-linear fit (variable slope, four parameters,calculated using GraphPad Prism v8.0.1) of the vitamin B_12_ yield with corresponding *r*² and EC_50_ values. Average vitamin B_12_ content in the fermentation medium ± S.D. is given for fermentations at 51 mM and 85 mM GB.

Using the Cob proteins of *S. meliloti* as references, the corresponding *cob* genes were annotated in the *E. adhaerens* LU2703 genome (Balabanova *et al*., 2021), (results are summarized in Table S1). In both strains, the majority of *cob* genes are organized in two large clusters (Fig. 4A), whose gene content broadly reflects the biochemical stages of vitamin B_12_ synthesis. The size of cluster 1 is 18.3 kbp in *E. adhaerens* LU2703 and 26.9 kbp in *S. meliloti* 102F34. It encodes, with the exception of *cobA* (the gene for the initial corrin ring methylase), exclusively genes whose products are required for the middle stage of cobalamin biosynthesis, which comprises cobalt insertion and amidation of hydrogenobyrinic acid. The cluster 1 size difference between the two strains is attributable to a locus within cluster 1 that contains different ORFs in each organism. None of these ORFs seem to be related to vitamin B_12_ biosynthesis (Fig. 4A). Cluster 2 is approximately 7.5 kbp in size in both organisms and encodes enzymes involved in the first stage of cobalamin biosynthesis. The CobA methylase is the only enzyme required for corrin ring methylation that is not encoded in cluster 2. Most of the enzymes encoded in cluster 2 have been shown to use SAM as their methyl group donor (White and Demain, 1971; Balabanova *et al*., 2021).

**Fig. 4:**
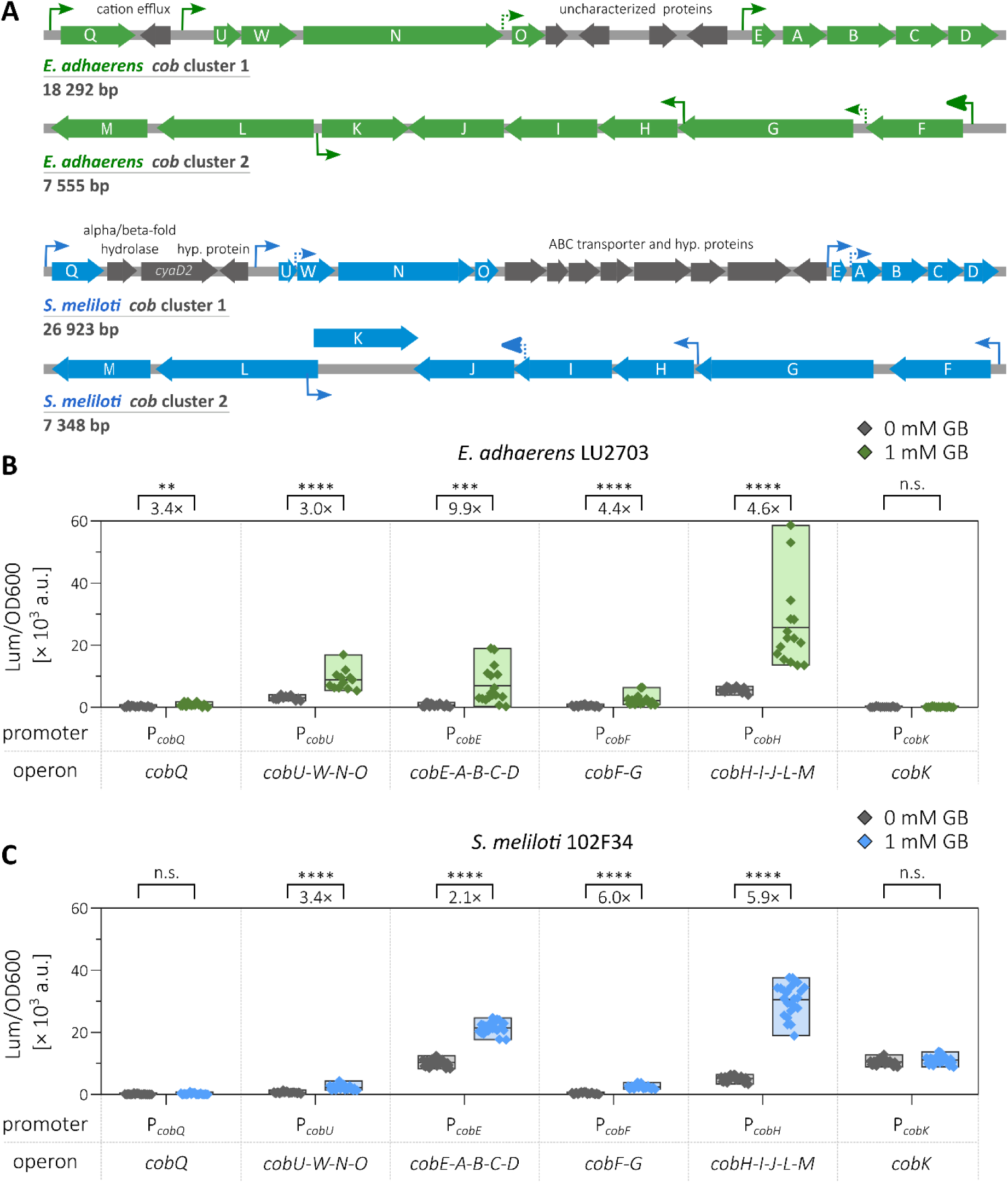
Activity of transcriptional-translational promoter probes in the *cob* biosynthetic gene cluster of *S. meliloti* and *E. adhaerens*. **(A)** Schematics of the two *cob* gene clusters of *E. adhaerens* LU2703 (top, green) and *S. meliloti* 102F34 (bottom, blue) and the location of promoters (arrows). Promoters outlined as dashes were suggested by Cappable-seq data but showed no activity in the reporter assay above background levels (data shown in Fig. S9). **(B, C)** Transcriptional-translational promoter-probe assay of regions upstream of *cob* genes with annotated promoters in *E. adhaerens* LU2703 (B) and *S. meliloti* 102F34 (C) after 6 h of cultivation in MOPS-buffered minimal medium with (1 mM) or without (0 mM) GB in microplates. Data based on 6 to 12 biological replicates measured twice on different days. Promoter fragments contained approximately 400 bp (or 800 bp for P*_cobU_*) upstream of the respective *cob* gene’s start codon. Statistical significance was calculated by multiple *t* test (n.s.: not significant, **: *p_adj_* < 0.001, ***: *p_adj_* < 0.000 1 ****: *p_adj_* < 0.000 01).

Using our Cappable-seq data for *E. adhaerens* LU2703, we confirmed that the *cob* genes are organized in multiple operons within their clusters (Fig. S8, Table S2), consistent with previously reported data for *S. meliloti* (Schlüter *et al*., 2013; Meier *et al*., 2024). Differences between *E. adhaerens* LU2703 and *S. meliloti* 102F34 were observed with regard to secondary TSS within *cob* cluster 1 and 2, suggested by the highly sensitive Cappable-seq analysis (dashed outlined promoters in Fig. 4A). However, we failed to demonstrate measurable promoter activities associated with these secondary TSS under our experimental conditions (Fig. S9).

Taken together, the genetic and transcriptional organization of the *cob* clusters appeared to be conserved between the vitamin B_12_ overproducer *E. adhaerens* LU2703 and the low-producer *S. meliloti* 102F34. Our TSS analysis determined the precise locations of the main promoters for *cobQ*, *cobUWNO*, *cobEABCD*, *cobHIJLM*, *cobFG*, and *cobK*, and thereby enabled investigating the effect of GB on the activity of these promoters.

### Transcriptional activity within rhizobial *cob* clusters is stimulated by GB

To investigate the transcriptional-translational effects of exogenously provided GB on *cob* gene expression, we generated promoter probe constructs corresponding to the TSS detected upstream of *cob* genes. Approximately 400 bp fragments upstream of the start codon of the TSS-associated coding region were fused to a *luxCDABE* reporter cassette on a single-copy plasmid and introduced to their cognate wild type, *E. adhaerens* LU2703 or *S. meliloti* 102F34. These reporter strains were grown in minimal medium with or without 1 mM GB. Luminescence and OD600 were measured after 6 h of incubation.

In *E. adhaerens* LU2703, the addition of 1 mM GB stimulated the activity of all measured *cob* promoters in clusters 1 and 2, except for the monocistronically counter-transcribed *cobK* gene (Fig. 4A, B). The *cobH* promoter exhibited the highest promoter activity in the presence of GB (4.6-fold induction). This promoter controls the *cobHIJLM* operon encoding four methyltransferases and one methylmutase responsible for corrin ring methylation. The strongest GB response was exhibited by the *cobE* promoter (9.9-fold induction), which controls the *cobEABCD* operon encoding the methyltransferase CobA that initiates the corrin ring methylation cascade.

In *S. meliloti* 102F34, the *cob* promoters responded to GB in a similar pattern. GB increased the relative luminescence 2.1- to 6.0-fold. The *cobH* and *cobE* promoters also showed the highest activities and were significantly stimulated in the presence of GB (5.9-fold and 2.1-fold, respectively). A notable difference between the two strains was the behavior of the *cobK* promoter, which is oriented opposite to the genes driven by the *cobH* promoter. The *cobK* gene encodes the precorrin-6A reductase, which catalyzes the NADPH-dependent reduction of precorrin-6A to precorrin-6B. In *E. adhaerens* LU2703, *cobK* promoter activity was generally low, bordering on the detection limit of the assay. In *S. meliloti* 102F34, its activity was intermediate between the stimulated and non-stimulated states of the *cobH* promoter, which controls a transcript encompassing *cobH*, *cobJ*, *cobL*, and *cobM*, as well as the antisense sequence to *cobK*.

We wanted to explore whether the regulation of the *E. adhaerens* LU2703 *cob* promoters was specific to the *Sinorhizobium*/*Ensifer* clade or had a phylogenetically broader effect. Therefore, we transferred luminescence reporter plasmids carrying *E. adhaerens* LU2703 *cobE*, *cobH*, and *cobU* promoters to *Rhizobium leguminosarum* Norway and *Agrobacterium tumefaciens* C58S (a spontaneous streptomycin-resistant mutant of the C58 strain). The resulting reporter strains were grown in minimal medium with and without 1 mM GB, and OD600 and luminescence was recorded (Fig. S10). Although overall promoter activities were low, we observed significant stimulation of the *cob* promoters in both *Rhizobiaceae* species in the presence of GB. This suggests that other members of the *Rhizobiaceae* family are also capable of GB-dependent stimulation of vitamin B_12_ biosynthesis.

GB-dependent stimulation of *cob* promoters was not only observed in the transcriptional-translational fusion reporter strains but also confirmed using strains carrying two sets of transcriptional promoter-reporter fusions with standardized 5’-UTRs. *E. adhaerens* LU2703 was equipped with plasmids carrying either long (between 220 bp and 315 bp upstream of the TSS) or short (100 bp upstream of the TSS) *cob* promoter fragments fused to the *luxCDABE* luminescence reporter operon (Fig. S11A). We then measured promoter activities of these transcriptional reporter strains cultured in minimal medium with and without 1 mM GB (Fig. S11B, C). All long transcriptional *cob* promoter-reporter fusions, covering the same upstream region as the fragments previously tested by transcriptional-translational fusions, showed significant GB-dependent stimulation (2.1 to 15.3 average fold induction). Among the short transcriptional fusions, only *cobE* and *cobQ* promoter fragments were sufficient for significant GB-dependent activation (2.4- and 1.7-fold, respectively).

To confine the DNA region responsible for GB-dependent regulation, we generated transcriptional fusion reporter plasmids in which successive parts of the *E. adhaerens* LU2703 *cobE* promoter were replaced with corresponding parts of the non-GB responsive promoter P*_MBHDPP_01747_* from the same strain (Fig. S12A). We introduced the resulting fusion plasmids into *E. adhaerens* LU2703 and measured promoter activities. Replacements of positions -100 to -61 abolished GB-dependent stimulation of the *cobE* promoter (Fig. S12B). Conversely, replacement of positions −100 to −41 of the P*_MBHDPP_01747_* promoter with the cognate sequence of the *E. adhaerens* LU2703 *cobE* promoter was sufficient to confer GB responsiveness (Fig. S12C). However, we were unable to identify a distinct and regulatory sequence motif within this fragment by promoter alignments (Fig. S13), motif search (Fig. S14), or palindrome search (Table S5).

### Vitamin B_12_ production rate correlates with GB-dependent stimulation of *cob* promoters in *E. adhaerens* LU2703

Having established that GB acts as an effector on *cob* promoter activity, we next examined whether this regulatory effect also occurs under vitamin B_12_ production conditions. We selected the *E. adhaerens* LU2703 *cobE* and *cobH* promoter fusions because both showed strong GB-dependent regulation and control operons within *cob* clusters 1 and 2. Together these operons encode the complete set of enzymes required for corrin ring modification, including all SAM-dependent methyltransferases (Fig. 4, Table S1). *E. adhaerens* LU2703 strains carrying *cobE* and *cobH* promoter-reporter plasmids were cultivated in batch fermentation using fermentation medium supplemented with increasing concentrations of GB. After five days of incubation, we measured vitamin B_12_ production by HPLC and promoter activities by determining relative luminescence (Fig. 5). The vitamin B_12_ production yields of the *cobE* and *cobH* promoter-reporter strains (Fig. 5A, D) closely mirrored that of *E. adhaerens* LU2703 wild type (Fig. 3), with both strains exhibiting the previously observed sigmoidal dependence on GB concentration. In both strains vitamin B_12_ yields reached saturation at 51 mM GB supplementation, as previously observed for the wild type, although absolute yields differed slightly. In parallel, both the *cobE* and *cobH* promoter activities responded to GB and followed sigmoidal dose-response curves (Fig. 5A, C). Half-maximal stimulation of the *cobE* promoter occurred at 35 mM GB, comparable to the GB concentration required for half-maximal vitamin B_12_ yield (27 to 32 mM GB; Fig. 3, 5B, D). The *cobH* promoter required less GB to be stimulated and reached half-maximal activity at 17 mM GB, whereas half-maximal vitamin B_12_ yields required higher GB concentrations. Repeating this experiment independently gave consistent results (Supplementary Data S1).

**Fig. 5:**
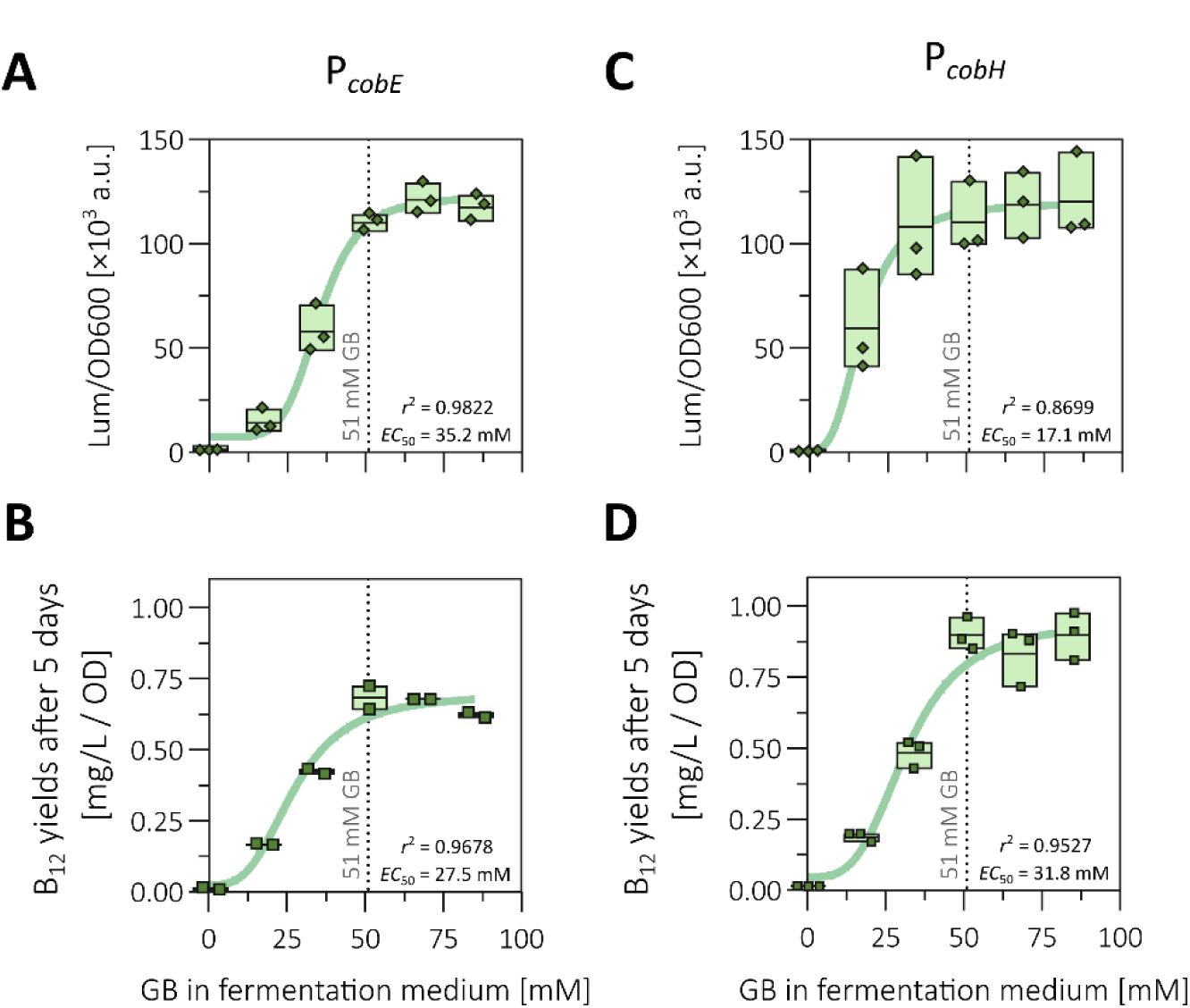
GB-dependent transcriptional stimulation of *cob* promoters correlates with productive vitamin B_12_ synthesis. *E. adhaerens* LU2703 carrying either *cobE* or *cobH* promoter-reporter plasmids (400 bp upstream of the start codon) were grown in MOPS-buffered fermentation medium with varying concentrations of GB in narrow-necked shaking flasks for five days. Activity of the *cobE* promoter **(A)** and the *cobH* promoter **(C)** was determined by diluting cultures after fermentation 1:100 and measuring luminescence and OD600 on a Tecan Infinite M Plex in microplates. Vitamin B_12_ yields are shown for strains carrying the *cobE* promoter-reporter plasmid **(B)** and the *cobH* promoter-reporter plasmid **(D)**. Green symbols represent biological replicates. The green line shows a non-linear fit (variable slope, four parameters, calculated using GraphPad Prism v8.0.1) with *r*^2^ and *EC*_50_ values given in each plot. The dashed line shows 51 mM GB, the concentration previously determined (Fig. 3) as sufficient for maximal vitamin B_12_ production.

Notably, strong promoter stimulation of the *cobE* and the *cobH* promoters during batch fermentation required roughly 50 times more GB compared to the promoter-reporter assays. As shown in Fig. S15, 1 mM GB, which had previously been sufficient for promoter stimulation in the promoter-reporter assays, was insufficient under production conditions to enhance either transcription levels or vitamin B_12_ synthesis. This discrepancy can be explained by the difference in cell densities between the two experimental setups. Batch fermentation in an optimized production medium over five days of incubation yields cell densities that are approximately 50- to 60-fold higher than those reached the in promoter-reporter assays conducted in 96-well microplates in minimal medium after 6 h of incubation (average OD600 of 4.9 versus 0.08 as measured in 96-well microplates). Thus, the proportionally higher GB requirement under fermentation conditions directly reflects the higher biomass present.

To investigate whether GB-dependent transcriptional stimulation of *cob* promoters is functionally important for vitamin B_12_ production, we chromosomally replaced the first 100 bp of the *E. adhaerens* LU2703 *cobE* promoter with either the non-GB-responsive P*_MBHDPP_01747_* promoter or the GB-responsive P*_MBHDPP_01747_*/P*_cobE_* hybrid promoter (Fig. S16). In previous measurements, P*_MBHDPP_01747_* exhibited the same activity level as P*_cobE_* in the absence of GB, regardless whether or not GB was present, while the P*_MBHDPP_01747_*/P*_cobE_* hybrid promoter exhibited a response to GB that was comparable to that or P*_cobE_* (Fig. S12). Replacement with the non-responsive promoter resulted in a 6.3-fold reduction in vitamin B_12_ yield after five days with 51 mM GB present in the growth medium (from 0.8 to 0.13 mg B_12_/L / OD600) (Fig. S16E). While wild type *E. adhaerens* LU2703 achieved a 20-fold higher yield in the presence of 51 mM GB than in its absence, the P*_MBHDPP_01747_*-mutant strain only achieved a 5.4-fold increase over its basal level. Both phenotypes were partially restored when the *cobE* promoter was replaced with the synthetic GB-responsive hybrid promoter (Fig. S16E). This strain yielded (0.42 ± 0.03) mg B_12_/L / OD600 corresponding to 52 % of the wild type level and achieved a 22-fold increase in B_12_ yield in the presence of GB compared to its absence. Taken together, these data demonstrate a tight coupling of GB availability to vitamin B_12_ production rate via transcriptional stimulation.

## Discussion

Industrial production of vitamin B_12_ relies exclusively on certain bacteria and archaea, with members of the order Hyphomicrobiales, particularly *E. adhaerens* and *S. meliloti,* emerging as natural overproducers following the aerobic biosynthetic route (Vu *et al*., 2013; Dong *et al*., 2016; Zhao *et al*., 2019; Bampidis *et al*., 2023; Liu *et al*., 2023). Here, we used the novel patent-restricted vitamin B_12_ high-producer strain *E. adhaerens* LU2703 to demonstrate that GB, beyond supplying methyl groups and biosynthetic precursors, also acts as a transcriptional effector on *cob* genes, in particular those encoding methyltransferases directly dependent on methyl group availability.

Vitamin B_12_ biosynthesis is energy- and resource-intensive, requiring the concerted action of 25 to 30 enzymes, and must be tightly regulated. Beyond the well-described vitamin B_12_-responsive riboswitches that feedback-regulate precursor uptake and biosynthetic genes (Rodionov *et al*., 2003), GB-dependent stimulation of vitamin B_12_ production has long been recognized (Demain *et al*., 1968)) and is industrially exploited by supplementing fermentation media with GB, for example via sugar beet molasses containing up to 6% (w/w) GB (Li *et al*., 2013).

Using promoter-reporter fusions in *E. adhaerens* LU2703 and *S. meliloti* 102F34, we show that GB stimulates the *cobE* and *cobH* promoters, which drive transcription of the two major *cob* operons encoding the methyltransferases responsible for sequential precorrin ring modifications from uroporphyrinogen III to hydrogenobyrinic acid. Promoter stimulation mirrored the GB-dependent increase in vitamin B_12_ yield during batch fermentation, with the *cobH* promoter activity reaching saturation before maximum production rates were achieved. This suggests that full production of the *cobH*-encoded enzyme is ensured prior to pathway saturation. This regulatory pattern is consistent with the dual metabolic role of GB, whose catabolism yields glycine as a direct precursor for 5-ALA synthesis and is thought to feed methyl groups into the SAM cycle required for tetrapyrrole ring methylations. GB-dependent stimulation of *cob* genes may thus represent a feed-forward mechanism that couples biosynthetic gene expression to the availability of a key metabolic substrate, thereby downregulating the pathway in the absence of GB as substrate. This function of GB in vitamin B_12_ biosynthesis requires its active catabolism. We observed GB-dependent activation of the degradation pathway promoters, most notably those controlling the demethylation steps that liberate methyl groups (P*_dmg2_*, P*_soxII_*). Together, this is consistent with a recent report showing that increased expression of the 5-ALA synthase *hemA* gene enhances vitamin B_12_ yield in a related *E. adhaerens* strain (Wang *et al*., 2025). A similar regulatory pattern is suggested for *P. denitrificans*, whose Cob proteins share high sequence similarity with those of *E. adhaerens* LU2703 (Table S6), and in which proteomic and metabolomic studies revealed increased abundance of vitamin B_12_ biosynthesis-related proteins and metabolites upon GB supplementation (Xia *et al*., 2015; Li *et al*., 2022).

While GB-dependent gene regulation has been described in *P. aeruginosa* and *B. subtilis* in the contexts of host colonization and osmoadaptation (Wargo *et al*., 2008; Nau-Wagner *et al*., 2012; Hampel *et al*., 2014), the promoters identified here showed no recognizable binding sites for GbdR-, GbsR- or OpcR-like regulators, and osmoadaptation did not appear to be the relevant context in either of the *Rhizobiaceae* species studied. The existence and identity of a putative GB-responsive transcriptional regulator therefore remains elusive. Biophysical mechanisms, such as GB-induced changes in DNA supercoiling or stabilization of protein-DNA complexes through its properties as a chemical chaperone (Kurz, 2008), may contribute, and warrant future investigation.

Understanding the regulatory principles that govern vitamin B_12_ biosynthesis is a prerequisite for rational strain engineering. The GB-dependent transcriptional stimulation of *cob* genes identified here represents a novel regulatory layer that can be exploited to optimize promoter design and fine-tune *cob* gene expression, thereby providing new molecular targets and strategies for the development of high-level vitamin B_12_ production strains.

## Experimental procedures

### Bacterial strains and media

*Escherichia coli* DH5α (Hanahan, 1983) was used for cloning and propagating plasmids. Consequently, each plasmid listed in Table S3 is associated with an *E. coli* DHα strain carrying that plasmid that was generated during this study. *E. coli* MT616 (Finan *et al*., 1986) was used as a helper strain for triparental conjugation. *E. coli* strains were cultivated in LB (5 g/L yeast extract, 10 g/L tryptone, 5 g/L NaCl) with appropriate antibiotics at 37 °C. *S. meliloti* 102F34 (Pichereau *et al*., 1998), *Agrobacterium tumefaciens* C58S (a spontaneously derived streptomycin-resistant C58S mutant provided by Antonio Lagares Jr.), *Rhizobium leguminosarum* Norway (Liang *et al*., 2018), and patent-restricted *E. adhaerens* LU2703 (DSM No. 34968), all members of the alphaproteobacterial family *Rhizobiaceae*, were used in this study. All alphaproteobacterial strains are listed in Table S3, and were cultivated in TY (3 g/L yeast extract, 5 g/L tryptone, 0.4 g/L CaCl_2_ · 2 H_2_O) or modified 3-(N-morpholino)propanesulfonic acid (MOPS)-buffered minimal medium (Zhan *et al*., 1991) (basic solution: 10 g/L MOPS, 10 g/L mannitol, 1.05 g/L (NH_4_)_2_SO_4_, 0.246 g/L MgSO_4_ · 7 H_2_O, adjusted to pH 7.2 using KOH, autoclaved; supplemented basic solution to a final concentration of 250 µM CaCl_2_, 10 mg/L FeCl_3_, 2 mM K_2_HPO_4_, 0.1 mg/L biotin, 0.1 mg/L thiamine-HCl, 0.1 mg/L pantothenate, 3 mg/L H_3_BO_3_, 2.23 mg/L MnSO_4_ · 4 H_2_O, 0.287 mg/L ZnSO_4_ · 4 H_2_O, 0.125 mg/L · CuSO_4_ · 5 H_2_O, 0.065 mg/L CoCl_2_ · 2 H_2_O, 0.12 g/L NaMoO_4_ · 2 H_2_O) at 30 °C. When indicated, the MOPS-buffered minimal medium contained alternative carbon sources to replace mannitol, maintaining equimolar carbon concentrations or no carbon source. Additionally, *E. adhaerens* LU2703 was cultivated in complex DIFCO Brain Heart Infusion medium (BD, Franklin Lakes, USA) supplemented with 2 g/L sucrose for determination of transcription start sites. Vitamin B_12_ fermentation was carried out in MOPS-buffered fermentation medium (“MOPS-B12 Fermentation Medium” and “MOPS-B12 Preculture Medium”, Table S4). Antibiotics were used whenever plasmids were introduced to any strain, specifically gentamycin (Gm, 10 µg/mL for *E. coli* DHα in liquid and on solid media, 20 µg/mL for *E. adhaerens* LU2703, *S. meliloti* 102F34, *A. tumefaciens* C58S, and *R. leguminosarum* Norway in liquid media, 30 µg/mL on solid media), spectinomycin (Sp, 100 µg/mL for *E. coli* DHα in liquid and on solid media, 200 µg/mL for *E. adhaerens* and *S. meliloti* both in liquid and on solid media), streptomycin (Str, 600 µg/mL for *E. adhaerens* LU2703, *S. meliloti* 102F34, *A. tumefaciens* C58S, and *R. leguminosarum* Norway in liquid and on solid media), hygromycin-B (Hyg, 100 µg/mL for *E. coli* DHα on solid media and 100 µg/mL for *E. adhaerens* LU2703 and *S. meliloti* 102F34 in liquid and on solid media) and chloramphenicol (Cml, 25 µg/mL for *E. coli* MT616 on solid media).

### Identification of homologous proteins

Basic local alignment search tool with protein sequence (BLASTp) analysis was performed on a local computer (Camacho *et al*., 2009) based on a database of all translated ORFs from *E. adhaerens* LU2703. The “blastp” (version 2.16.0+) command was run with default parameters. Characterized *S. meliloti* 102F34 proteins with presumed functions were used as queries. The gene related to the best result for each query was considered as a putative homolog from the *E. adhaerens* LU2703 genome.

### Plasmid construction and DNA oligonucleotides

All plasmids and details of their construction are listed in Table S3. DNA oligonucleotides for PCR reactions and sequencing can be found in Table S3. General techniques for cloning employed GoldenGate cloning following the MoClo standard and its extensions (Weber *et al*., 2011; Werner *et al*., 2012; Meier *et al*., 2024) as well as standard restriction-ligation cloning. Plasmids were introduced to *E. coli* DHα following established methods of transformation of chemically competent cells (Inoue *et al*., 1990).

### Transfer of DNA to Alphaproteobacteria

DNA was transferred to *E. adhaerens* LU2703, *S. meliloti* 102F34, *A. tumefaciens* C58S, and *R. leguminosarum* Norway by triparental conjugation. Briefly, alphaproteobacterial recipient strain, *E. coli* DH5α donor, and *E. coli* MT616 helper strain were streaked on TY or LB agar media with the respective antibiotics. Then, they were mixed in liquid TY without antibiotics in an approximately ratio of 3:1:1 (recipient:donor:helper). From that mixture, 10 to 50 µL were spotted on TY agar medium without antibiotics. Mating spots were incubated overnight at 30 °C. Afterwards, mating spots were dissolved in 500 µL TY and spread in serial dilutions on TY agar medium containing antibiotics appropriate for plasmid maintenance as well as streptomycin for counter-selecting the *E. coli* strains. Visible colonies appeared after 3 to 4 days of incubation at 30 °C. Biological replicates were individual single colonies from one conjugation event.

### Chromosomal manipulation

Marker-free changes to the *E. adhaerens* LU2703 chromosome were performed by double-homologous recombination using the homing endonuclease I-SceI as a counter-selectable marker (Viret, 1993). Briefly, an I-SceI recognition site (5’ TAGGGATAACAGGGTAAT 3’) together with two homologous regions (between 800 and 1000 bp each) flanking a cargo (if present) were cloned on a pS18mob2 plasmid (Kirchner and Tauch, 2003) or derivative thereof using standard procedure. Detailed constructions are outlined in Table S3. Plasmids were introduced to *E. adhaerens* LU2703, and integration was confirmed by growth on TY with spectinomycin. Single colonies were restreaked on TY with spectinomycin and afterwards allowed to grow in liquid TY without antibiotic selection pressure for 2 days at 30 °C, 200 rpm. A plasmid that carried the *I-sceI* gene under the control of an IPTG-inducible promoter as well as an *mcherry* reporter cassette was introduced by conjugation to single-integration mutants derived from multiple colonies. Mating spots were plated on TY with Str, Gm, and isopropyl β-D-1-thiogalactopyranoside (IPTG). Double homologous recombination mutants were identified by lack of growth on TY with Str and Sp, and confirmed by colony PCR and/or Sanger sequencing of the respective locus. Loss of the *I-sceI* plasmid was verified by lack of *mcherry*-derived fluorescence as measured with an Amersham ImageQuant 800 with an excitation wavelength at 535 nm. Biological replicates of marker-free mutants originated from individual colonies from the first homologous recombination.

### Isolation of genomic DNA

Genomic DNA was isolated from *E. adhaerens* LU2703 cells cultivated to a lawn on TY with streptomycin. Cells were harvested by centrifugation at 20 000 rcf for 10 min. DNA was isolated from cell pellets with the NucleoSpin Microbial DNA Mini-Kit (Macherey-Nagel, Düren, Germany) according to the manufacturer’s instructions.

### Whole genome sequencing

Whole genome sequencing of *E. adhaerens* LU2703 was performed via a commercial service (Plasmidsaurus, Cologne, Germany) using amplification-free long-read Oxford Nanopore sequencing (BioProject PRJNA1505600; accession number: JCBRLB000000000). A total read count of 138 345 (longest read: 98 590 bp) was reached, resulting in a coverage of 78.9. The assembly consisted of three replicons, named scaffold1, scaffold2, and scaffold3.

### Purification of total RNA

For total RNA purification, cultures were grown and 0.5 mL culture broth was harvested by centrifugation at 12 000 rcf at 4 °C for 1 min. After removal of the supernatant, pellets were frozen in liquid nitrogen and stored at -80 °C until RNA extraction. RNA isolation was performed using the RNeasy Mini Kit (QIAGEN, Venlo, Netherlands) with on-column DNaseI digest according to the manufacturer’s instructions. RNA was eluted twice with 30 µL RNase-free water and stored at -80 °C until further usage. Concentrations were measured using a NanoDrop 2000 (Thermo Fisher Scientific, Waltham, USA), and RNA integrity was validated by agarose gel electrophoresis and analysis on a Bioanalyzer (Agilent Technologies, Santa Clara, USA).

### Determination of transcription start sites

For determination of transcription start sites (TSS), *E. adhaerens* LU2703 was cultured in 50 mL complex DIFCO Brain Heart Infusion medium (BD, Franklin Lakes, USA) supplemented with 2 g/L sucrose at 30 °C, 200 rpm shaking. Samples were taken at early exponential phase (32 h, OD600 ∼1) and mid exponential phase (38 h, OD600 ∼6). From 3 replicate cultures, 2 samples of 0.5 mL were taken at each time point. For RNA extraction, all samples from the two time points were pooled (T_32h_ rRNA ration (23S/16S): 1.5, RIN (RNA Integrity Number): 10; T_38h_ rRNA ratio (23S/16S): 1.5, RIN: 9.9). Both pools were mixed in a 1:1 ratio, Cappable-seq (Ettwiller *et al*., 2016) was carried out by a commercial service (Vertis Biotechnologie AG) (BioProject PRJNA1506472; SRA accession number: SRR39971765). Cappable-seq reads were mapped to the *E. adhaerens* LU2703 genome with CLC Genomics Workbench (v25.0.3, QIAGEN Aarhus A/S, Aarhus, Denmark) using “Map Reads to Reference” function with default parameters (Match Score: 1, Mismatch cost: 2, Linear gap cost, Length fractions: 0.5, Similarity fraction: 0.8). 5’ ends of mapped reads were considered putative TSS when at least *n* = 30 read ends mapped to a given position.

### Transcriptional and translational promoter-probe assays

Activity of promoter sequences was quantified by fusing respective DNA fragments to the *luxCDABE* luminescence-generating operon from *Photorhabdus luminescens*. Measurements were performed as previously established (Meier *et al*., 2024), unless stated otherwise (Fig. 5, S15). In brief, *E. adhaerens* LU2703, *S. meliloti* 102F34, *A. tumefaciens* C58S, or *R. leguminosarum* strains carrying luminescence-reporter cassettes on single-copy pABC plasmids (Döhlemann *et al*., 2017; Meier *et al*., 2024) were inoculated in TY with appropriate antibiotics in 96-well clear Greiner-F plates sealed with non-breathable seal and incubated at 30 °C, 1200 rpm on a Heidolph Titramax 1000 shaker. After 1:10 dilution, MOPS-buffered minimal medium in fresh plates were inoculated and incubated overnight as before. Assays were carried out employing a robotic platform (including an Tecan Evo Freedom 200 automated liquid handling system, a LiCONiC StoreX series incubator, and a Tecan Infinite M200 Pro Reader) and using black 96-well microplates (µCLEAR Black, F-Bottom, Greiner Bio-One, Frickenhausen, Germany) and lids with condensation rings for cell cultivation in MOPS-buffered minimal medium. Cell cultures were synchronized to an OD600 of 0.05, then GB, choline or water was added followed by incubation at 30 °C for 12 h, shaking vigorously every 1 h. Measurements were taken at timepoints 0, 3, 6, 9, and 12 h. Relative luminescence was calculated by dividing raw luminescence values (integration time: 250 ms) by blank-corrected OD600 for each respective well. OD600 or relative luminescence values below 0 usually indicated lack of growth or resulted from measurement errors and were therefore set to 0 during evaluation.

### Production of vitamin B_12_

Microbial production of vitamin B_12_ by *E. adhaerens* LU2703 and *S. meliloti* 102F34 was performed in narrow-necked shaking flasks under microoxic conditions. Fermentation took place in a MOPS-buffered fermentation medium (“MOPS-B_12_”). All components for the fermentation preculture and fermentation main culture media are listed in Table S4. Strains were streaked on TY agar medium (with respective antibiotics whenever plasmid maintenance was necessary) and incubated overnight at 30 °C. Subsequently, 10 mL fermentation preculture medium were inoculated in shaking flasks and incubated overnight at 30 °C, 200 rpm in a New Brunswick Scientific Innova 44 incubator (Edison, New Jersey, United States). Precultures were then adjusted to an OD600 of 0.2 using preculture medium and 4 mL of the adjusted preculture was added to each 20 mL fermentation medium in narrow-necked shaking flasks. The final fermentation cultures usually contained 51 mM GB. For experiments that used different GB concentrations, these are specified in the respective figure captions (Fig. 3, 5, S15). Narrow-neck shaking flasks were covered with cloth to allow minimal gas exchange and create microoxic fermentation conditions. After 5 days of growth at 30 °C and 200 rpm (New Brunswick Innova 44), multiple aliquots of 500 µL per sample were collected and frozen at -20 °C until further processing. Replicates for each experiment were either biological replicates from a conjugational event or three independent pre-cultures of wild type strains.

### Quantification of vitamin B_12_

Vitamin B_12_ occurs in multiple isoforms (e.g. hydroxy cobalamin, adenosyl cobalamin). Prior to quantification, all vitamin B_12_ samples were treated with NaCN to convert all cobalamins into cyanocobalamin. 500 µL fermentation culture were mixed with 500 µL water and 24 µL 10% NaCN and heated at 80 °C for 30 min after which samples were allowed to cool at room temperature for 10 min before filtering with a pore size of 0.22 µm. Vitamin B_12_ samples were analyzed on a 1260 Infinity HPLC system (Agilent Technologies, Waldbronn, Germany) using an Aqua® LC Column (3 µm C18 125 Å, 150 x 4.6 mm, Ea; Phenomenex LTD, Aschaffenburg, Germany) with a SecurityGuard (AQ C18 column with 3.2 to 8.0mm internal diameters; Phenomenex LTD, Aschaffenburg, Germany) and UV/vis detection at 240 nm (BW4) and 360 nm (BW40). A mobile phase gradient at 40 °C with 0.1 % phosphoric acid (eluent A) and acetonitrile (eluent B) was used as follows: 0 min (10% B), 14 min (22 % B), 20 min (100 % B), 22 min (100 % B). The analysis was calibrated with cyanocobalamin standards which were synthesized via cyanide conversion of commercially available adenosyl cobalamin.

### Search for potential DNA motifs

Sequences of promoter probes that were stimulated by GB were subjected to motif discovery search using the MEME suite (Bailey *et al*., 2015). Sequences were scanned for motifs using Multiple Em for Motif Elicitation (MEME) v5.5.9 (Bailey and Elkan, 1994) with default parameters. Additionally, promoter sequences were screened for enrichment of known bacterial transcriptional regulator binding sites using Simple Enrichment Analysis (SEA) v5.5.9 (Bailey and Grant, 2021) with default parameters.

### Statistical methods

Statistical significance was calculated with Graphpad Prism v8.0.1 employing the multiple *t* test with corrections using the Holm-Šidák method. Fits to dose-response curves were performed with Graphpad Prism v8.0.1 using the “Non-linear Fit” function (variable slope, four parameters).

When measurement results are referenced in the text, they are presented as the average result ± standard deviation (S.D.) based on at least three replicate measurements.

## Supporting information

Supplemental Data 1, numerical values for figures in main manuscript

Supplemental Data 1, numerical values for supplemental figures

## Author Contribution

The study was designed and conceived by CR, TH, KS, HS, OZ, and AB. Experiments were conducted and evaluated by CR, MW, TH, and PC. JG conducted experiments. FU performed genome sequencing experiments. RNA sequencing experiments were performed by PC and data was processed by CR. All authors were involved in drafting the manuscript.

## Acknowledgements

We thank Prof. Erhard Bremer and Dr. Antonio Largares Jr. for providing *S. meliloti* 102F34 and *Agrobacterium tumefaciens* C58S, respectively.

## Data availability

The data that supports the findings of this study are available in the supplementary material of this article (Supplementary Data S1 and S2). *E. adhaerens* LU2703 genome sequencing (BioProject PRJNA1505600) was deposited at GenBank (accession number: JCBRLB000000000) and Cappable-seq data (BioProject PRJNA1506472) generated in this study was deposited in the Sequence Read Archive (SRA) (accession number: SRR39971765).

