## Supplemental Figures S1 to S16 for "Glycine betaine-mediated transcriptional stimulation links its catabolism and vitamin B12 biosynthesis in *Rhizobiaceae*"


The following supplementary information is available as separate files:

- **Table S1:** Comparison of protein sequences involved in the uptake, synthesis and catabolism of glycine betaine and the synthesis of vitamin $B_{12}$ between *S. meliloti* 102F34 and *E. adhaerens* LU2703
- **Table S2:** Transcription start sites derived from Cappable-seq analysis of *E. adhaerens* LU2703
- **Table S3:** Plasmids (A), oligo nucleotides (B), and rhizobial strains (C) used in this study
- **Table S4:** Media and solutions used for the fermentation of vitamin $B_{12}$
- **Table S5:** Identification of palindromes in -100/-41 positions of GB-regulated promoters from *S. meliloti* 102F34 and *E. adhaerens* LU2703
- **Table S6:** Comparison of protein sequences involved in the synthesis of vitamin $B_{12}$ between *E. adhaerens* LU2703 and *Pseudomonas denitrificans*

- **Data S1:** Data used to generate main figures
- **Data S2:** Data used to generate supplementary figures

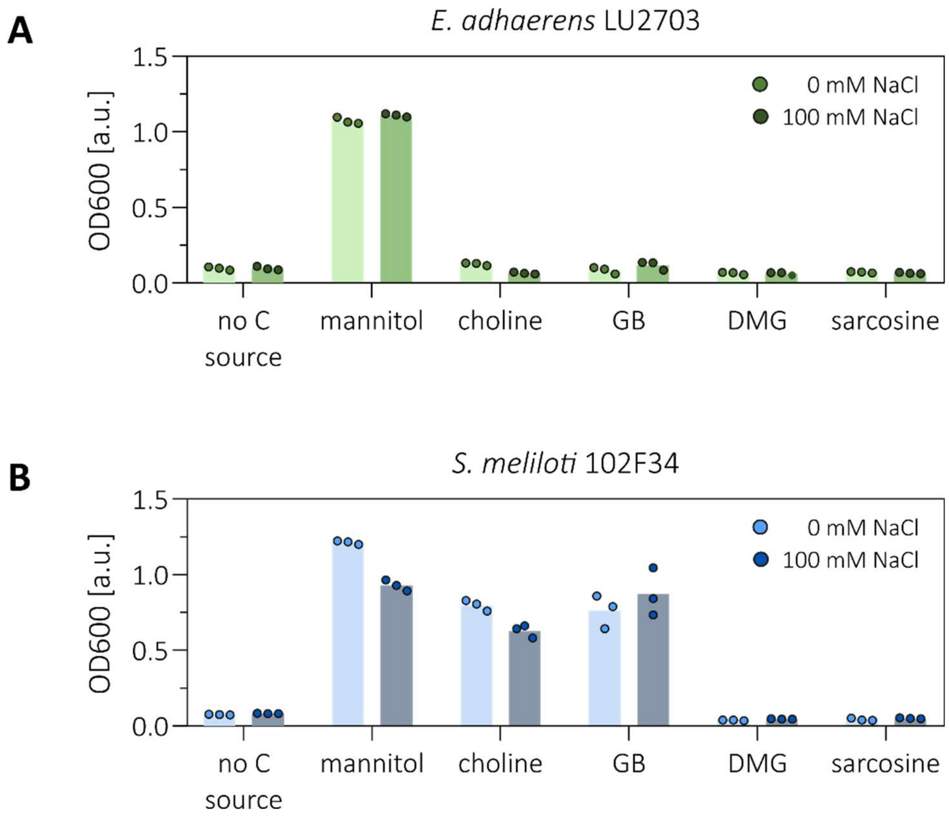


**Fig S1: Ability of *E. adhaerens* LU2703 and *S. meliloti* 102F34 to use substances related to glycine betaine as sole carbon sources in the presence or absence of NaCl.** Growth of *E. adhaerens* LU2703 (**A**, green bars and circles) and *S. meliloti* 102F34 (**B**, blue bars and circles) and after 48 h in MOPS-buffered minimal medium using either mannitol, GB, or metabolic precursors and products of GB as sole carbon source. Media were either prepared without additional NaCl or supplemented with 100 mM NaCl to introduce mild osmotic stress. Cultures were grown in microplates and measured on a Tecan Infinite M Plex reader. Precultures were grown in TY medium supplemented with 600 µg/mL streptomycin and washed twice with minimal medium without carbon source before inoculation.

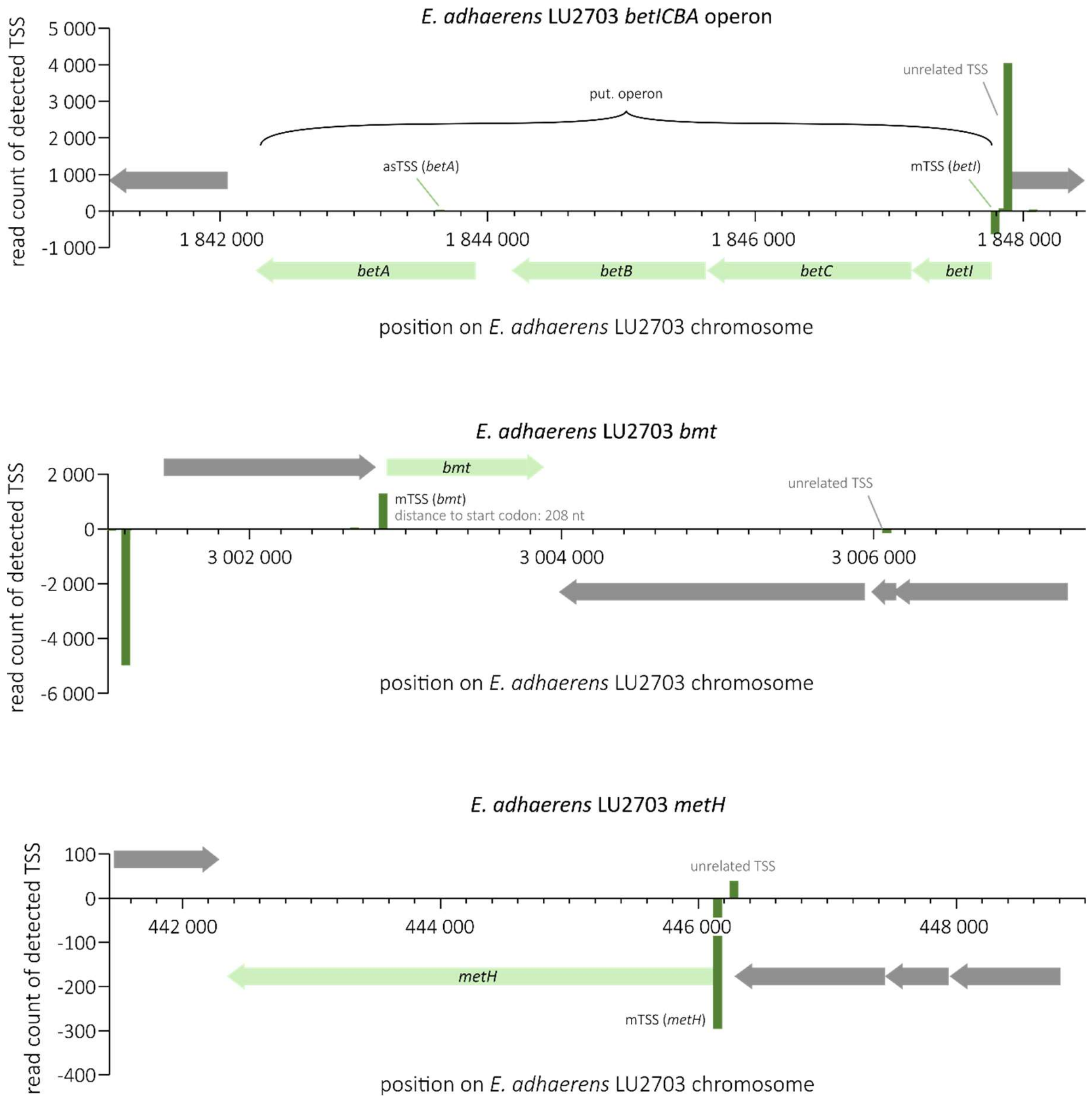


**Fig. S2: Transcription start sites for *E. adhaerens* LU2703 *betICBA* operon, *bmt*, and *metH*.** Arrows indicate genes, with light green highlighting relevant genes and grey indicating unrelated genes. Green bars indicate the number of detected TSS at a specific position either in the (+) strand (positive values) or the (-) strand (negative values). mTSS: TSS likely corresponding to an mRNA. asTSS: TSS likely corresponding to an anti-sense RNA.

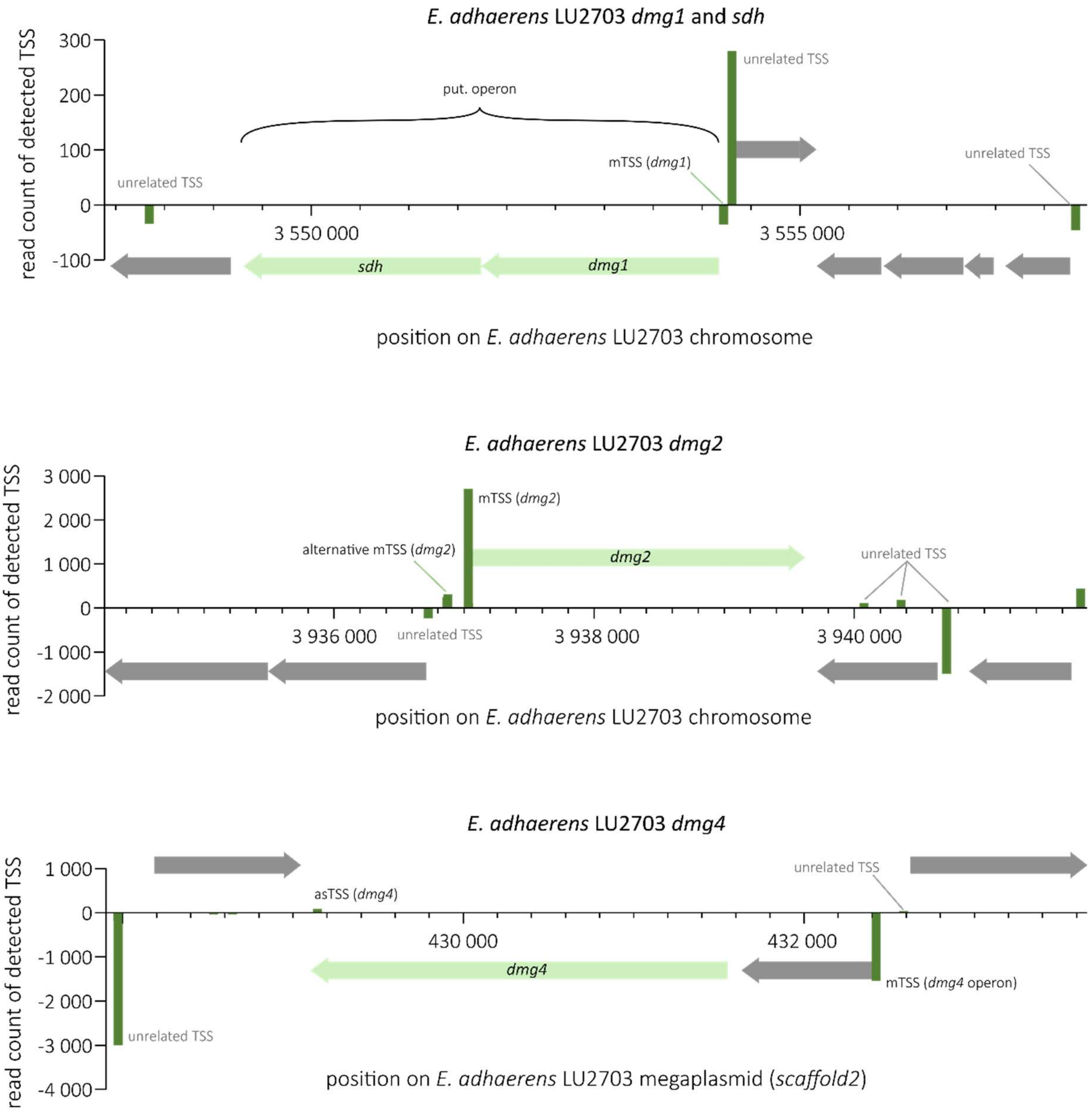


**Fig. S3: Transcription start sites for *E. adhaerens* LU2703 *dmg1*, *sdh*, *dmg2*, and *dmg4*.** Arrows indicate genes, with light green highlighting relevant genes and grey indicating unrelated genes. Green bars indicate the number of detected TSS at a specific position either in the (+) strand (positive values) or the (-) strand (negative values). mTSS: TSS likely corresponding to an mRNA. asTSS: TSS likely corresponding to an anti-sense RNA.

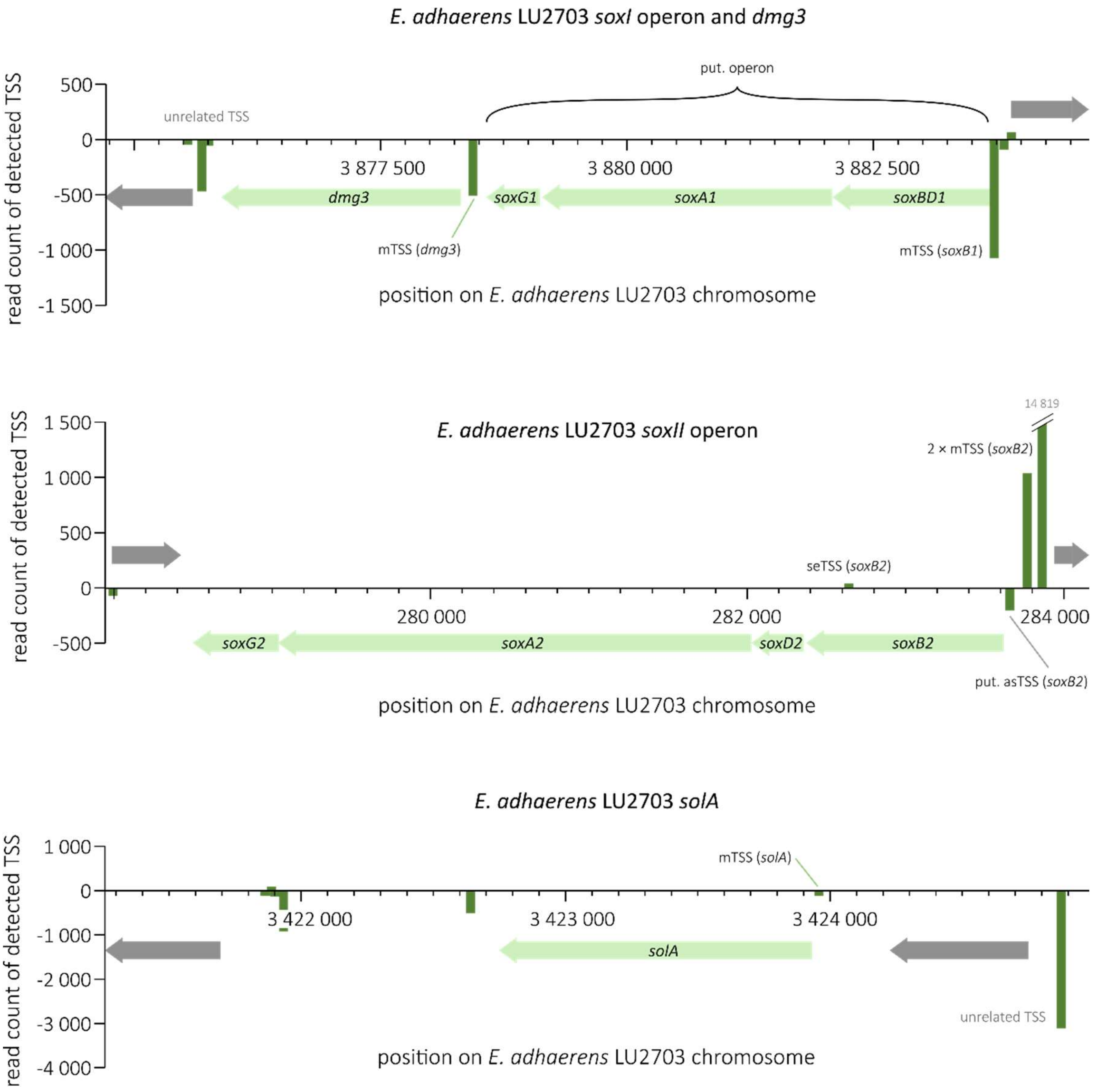


**Fig. S4: Transcription start sites for *E. adhaerens* LU2703 *soxI* operon, *dmg3*, *soxII* operon, and *solA*.** Arrows indicate genes, with light green highlighting relevant genes and grey indicating unrelated genes. Green bars indicate the number of detected TSS at a specific position either in the (+) strand (positive values) or the (-) strand (negative values). mTSS: TSS likely corresponding to an mRNA. asTSS: TSS likely corresponding to an anti-sense RNA. seTSS: TSS likely corresponding to a sense RNA.

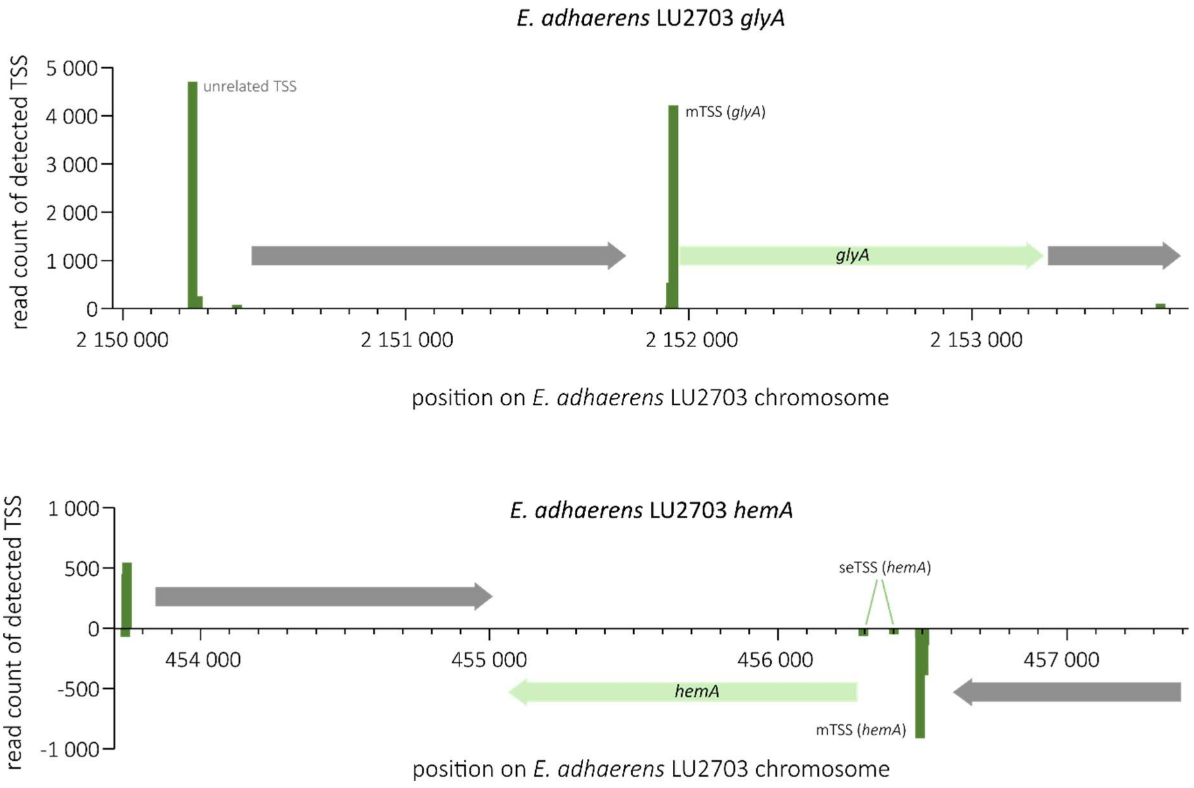


**Fig. S5: Transcription start sites for *E. adhaerens* LU2703 *glyA* and *hemA*.** Arrows indicate genes, with light green highlighting relevant genes and grey indicating unrelated genes. Green bars indicate the number of detected TSS at a specific position either in the (+) strand (positive values) or the (-) strand (negative values). mTSS: TSS likely corresponding to an mRNA. seTSS: TSS likely corresponding to a sense RNA.

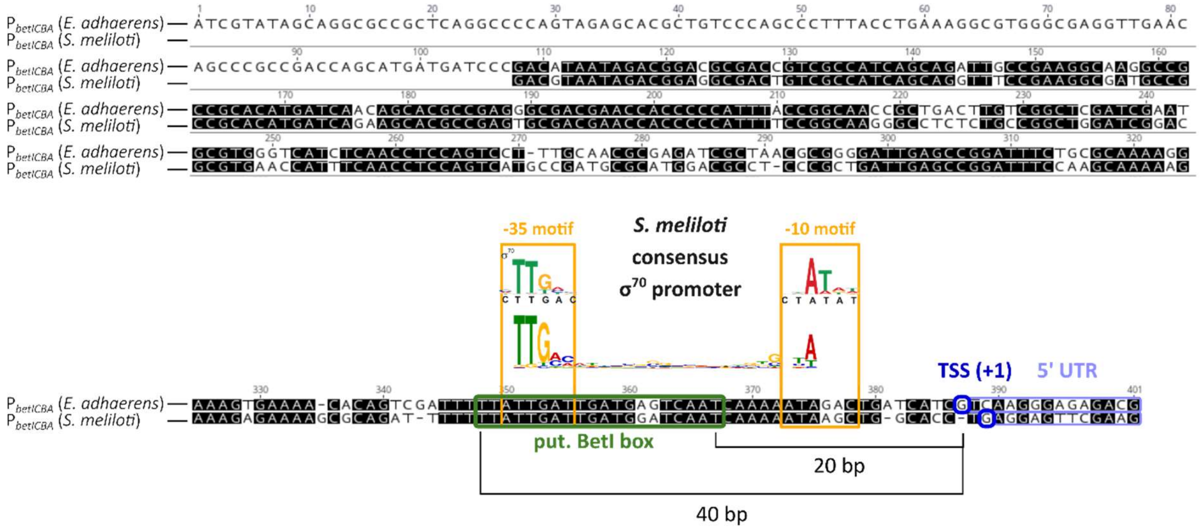


**Fig. S6: Alignment of the first 400 bp and 290 bp upstream of the *betICBA* operon from *E. adhaerens* LU2703 and *S. meliloti* 102F34, respectively, that were used as promoter probes.** Green box shows putative binding sites for BetI, the transcriptional TetR-type repressor of the *betICBA* operon. This binding site is conserved between both species except a 2-nt indel. Blue boxes show the transcription start site (TSS, +1) of each species. Given the gaps in the alignment, the putative BetI box is equidistant from both the *E. adhaerens* as well as the *S. meliloti* TSS (20 bp to 40 bp). Yellow boxes show the consensus promoter motifs of $\sigma^{70}$-dependent promoter from *S. meliloti*. Light blue boxes mark the 5' UTR.

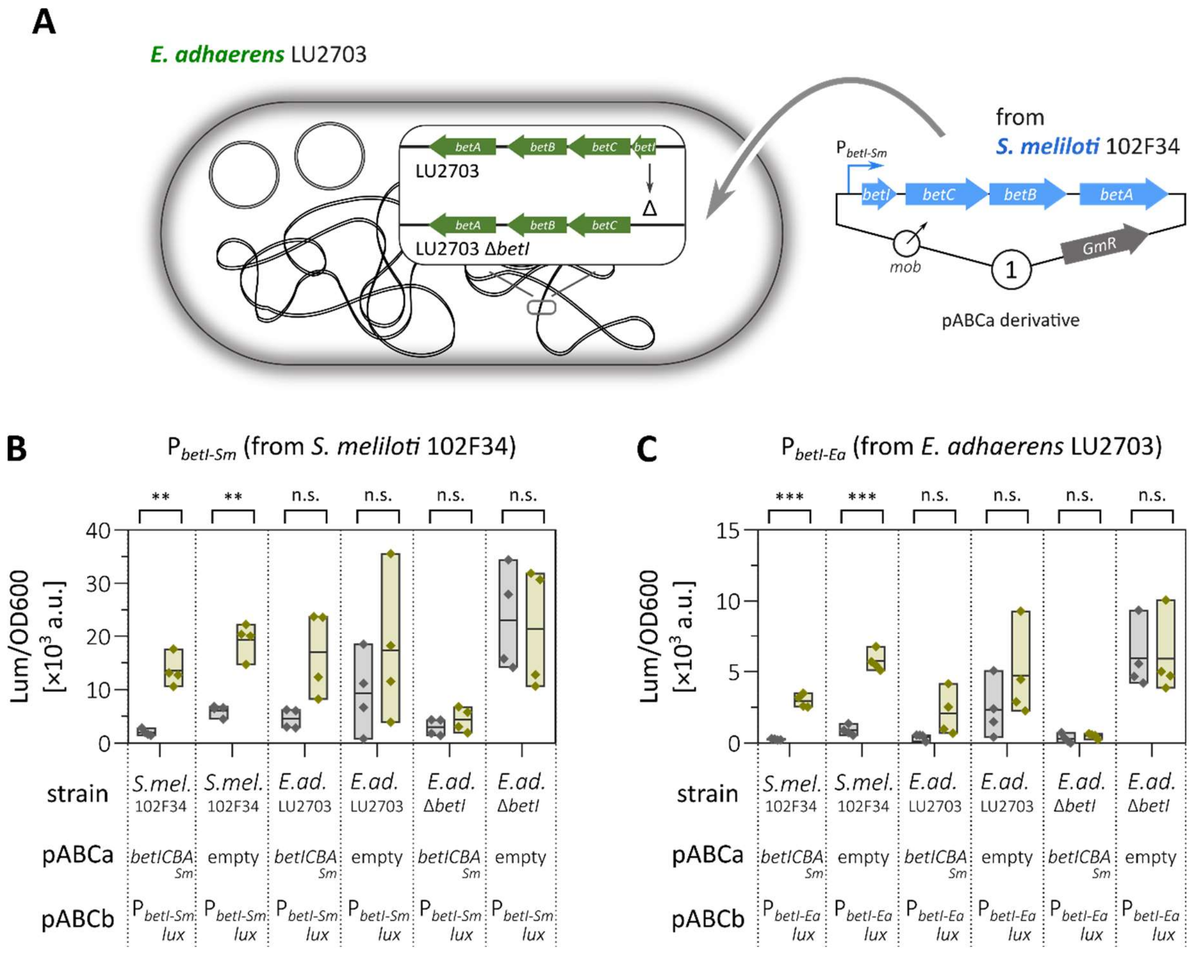


**Fig. S7: Characterization of *betI* promoters from *S. meliloti* 102F34 and *E. adhaerens* LU2703. (A)** *S. meliloti* 102F34 and *E. adhaerens* LU2703 (both wild type and *betI* deletion mutant) were equipped with an additional copy of the $P_{betI}$-*betICBA* operon from *S. meliloti* 102F34 or an empty plasmid. Transcriptional-translational $P_{betI}$-*luxCDABE* reporter fusions were introduced on another single-copy number plasmid. **(B, C)** Activity of the *S. meliloti*-derived $P_{betI\text{-}Sm}$ (B) or the *E. adhaerens*-derived $P_{betI\text{-}Ea}$ (C) promoter fragments 12 h after addition or not of 1 mM choline in MOPS-buffered minimal medium. No significant difference was detected in *E. adhaerens* strains upon addition of choline, regardless of promoter or presence of *betI* fragments. Significance was calculated using multiple *t* test (n.s.: not significant, **: $p_{adj} < 0.001$, ***: $p_{adj} < 0.000\,1$).

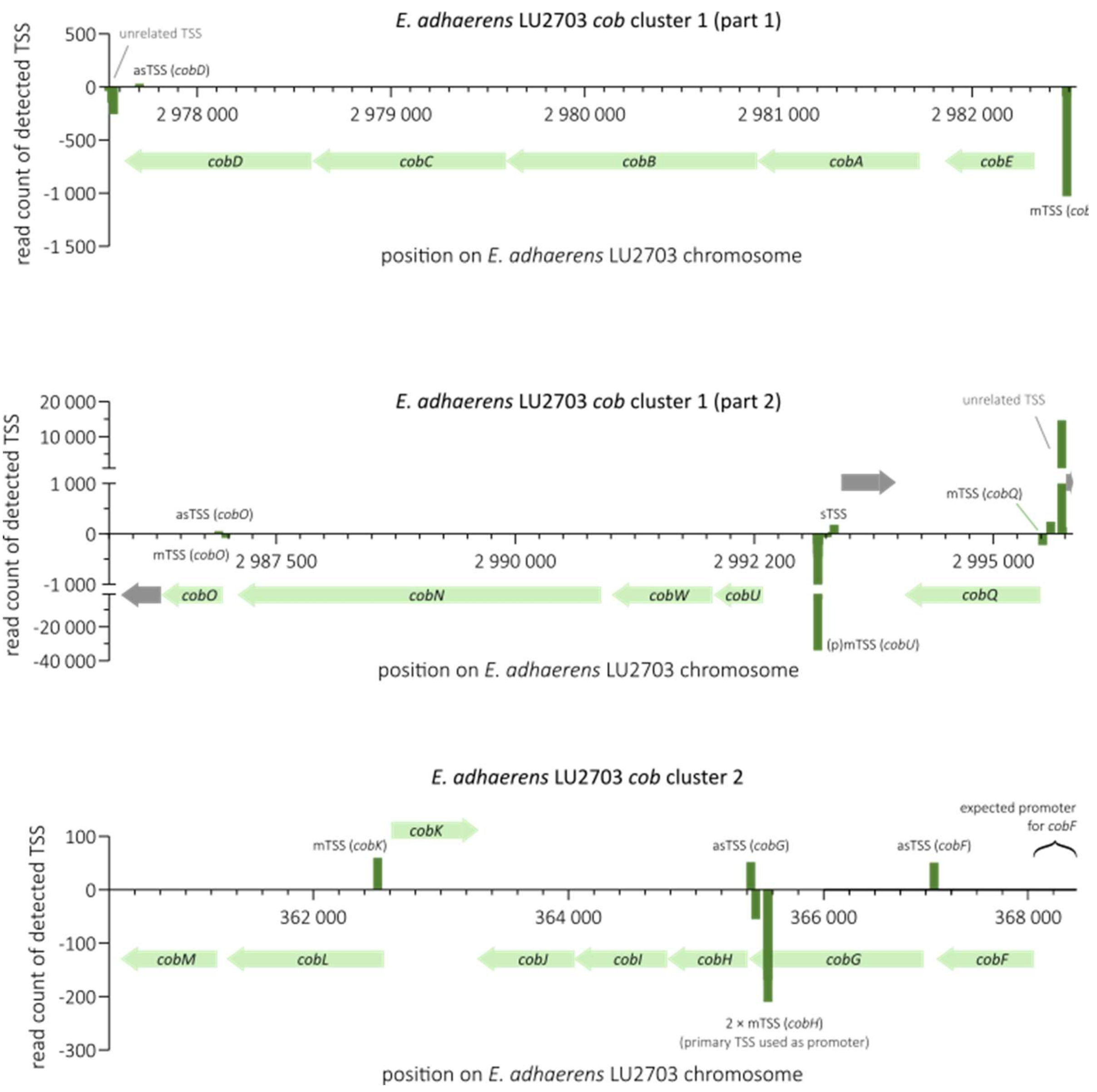


**Fig. S8: Transcription start sites for *E. adhaerens* LU2703 *cob* clusters 1 and 2.** Arrows indicate genes, with light green highlighting relevant genes and grey indicating unrelated genes. Green bars indicate the number of detected TSS at a specific position either in the (+) strand (positive values) or the (-) strand (negative values). mTSS: TSS likely corresponding to an mRNA.

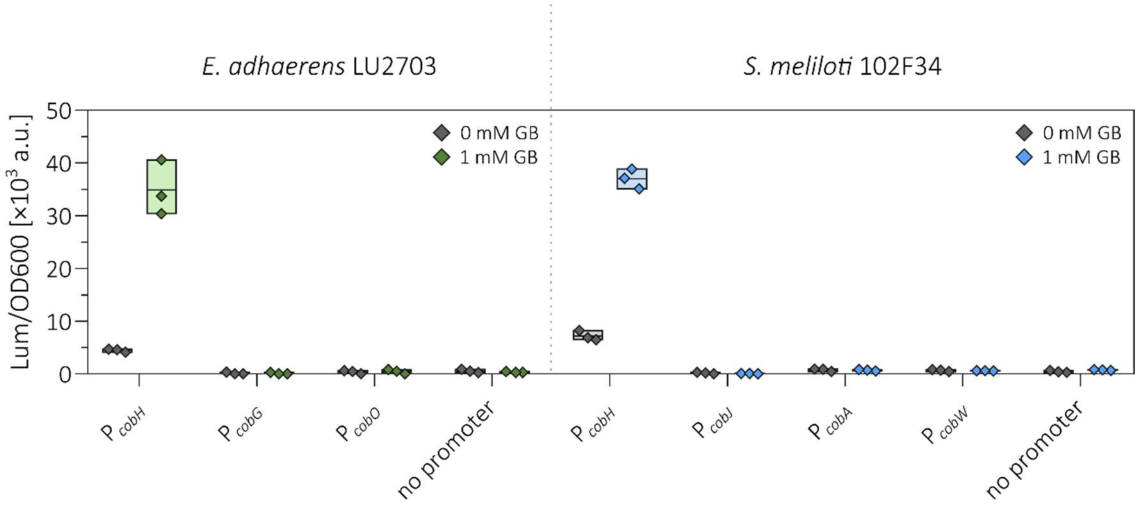


**Fig. S9: Not all TSS suggested by Cappable-seq transcriptome analysis of cells grown in minimal medium reflect promoters.** Promoter probes were constructed for sequences upstream of *cobG* and *cobO* from *E. adhaerens* LU2703 and upstream of *cobJ*, *cobA*, and *cobW* from *S. meliloti* 102F34 analogous to promoter probes described in Fig. 4. Strains carrying transcriptional-translational promoter-*luxCDABE* reporter constructs on a single-copy number plasmid were cultured in MOPS-buffered minimal medium with 0 mM or 1 mM GB for 6 h in microplates before data was taken with a Tecan Infinite M200 Pro reader. While control promoter $P_{cobH}$ showed strong stimulation and measurable activity even without GB for both species, all other promoters remained on levels comparable to control strains that carried an empty vector ("no promoter").

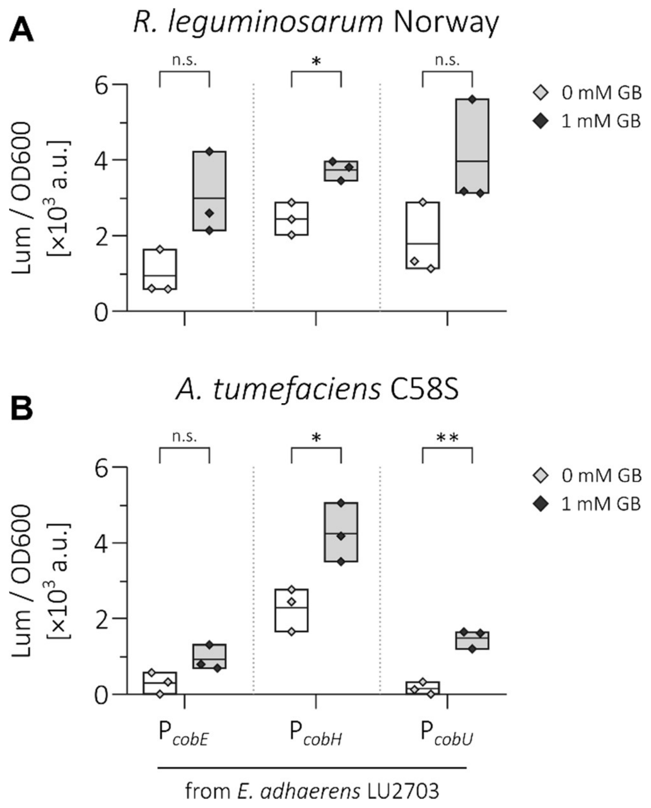


**Fig. S10: Activity of *cob* promoter-reporter fusions from *E. adhaerens* LU2703 in the presence and absence of 1 mM GB in *Rhizobium leguminosarum* Norway (A) and *Agrobacterium tumefaciens* C58S (B).** Strains were equipped with transcriptional-translational promoter probes on a single-copy number plasmid and were grown in MOPS-buffered minimal medium supplemented with either 0 or 1 mM GB. Data based on 3 biological replicates and taken after 15 h (for *A. tumefaciens* C58S) or 18 h (for *R. leguminosarum* Norway). Statistical significance was calculated by multiple *t* test (n.s.: not significant, *: $p_{adj} < 0.05$, **: $p_{adj} < 0.001$).

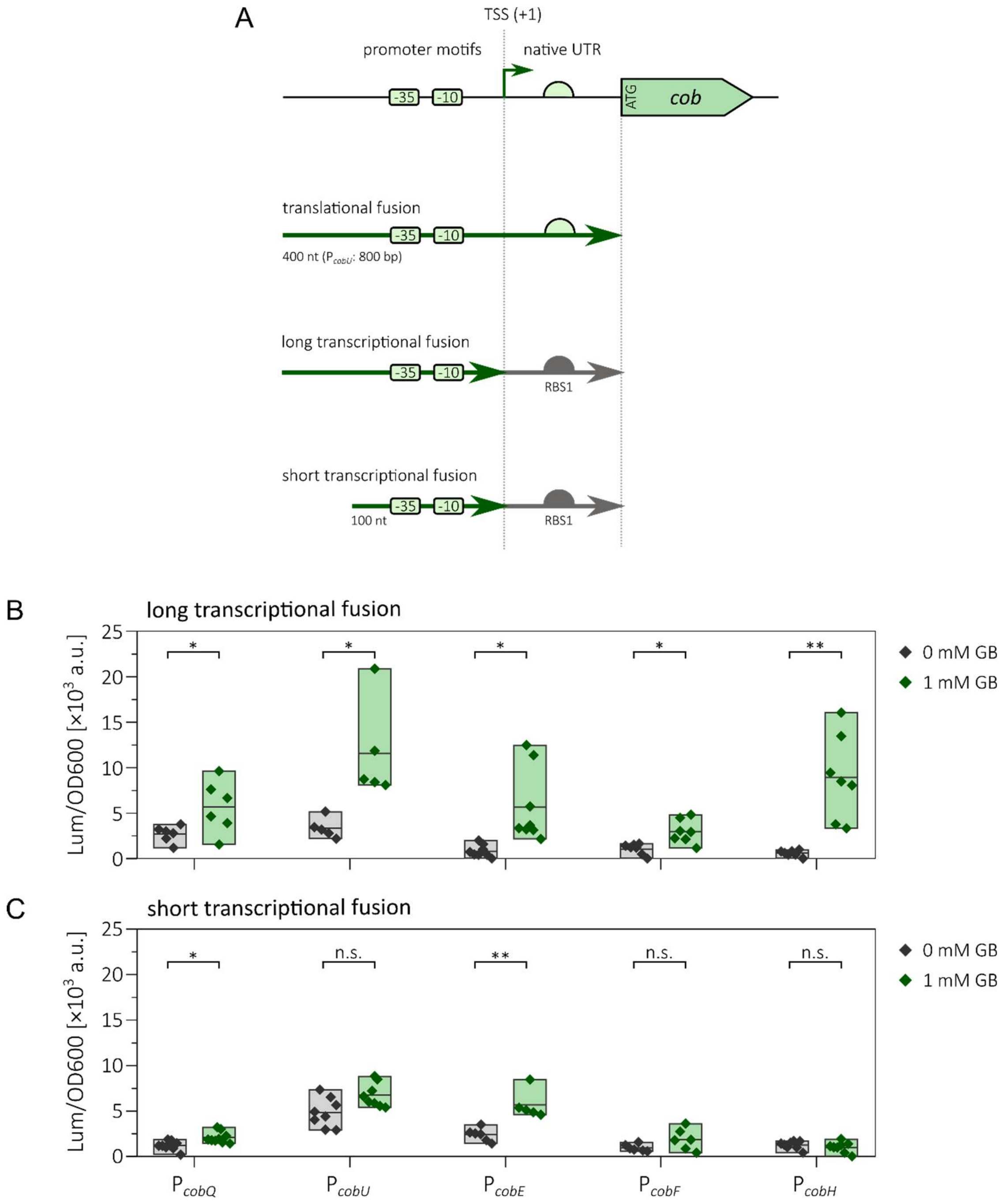


**Fig. S11: Stimulation of *E.adhaerens* LU2703 *cob* transcriptional fusion reporters by GB. (A)** Design of transcriptional-translational and transcriptional probes that were characterized by fusion to the *luxCDABE* reporter operon on single-copy number plasmids. Transcriptional-translational fusions were designed by cloning the first ca. 400 bp (800 bp in the case of *cobU*) upstream of the start codon of the respective *cob* CDS directly to the reporter cassette. Activity of these types of constructs is depicted in Fig. 4. Two types of transcriptional fusion reporters were evaluated: long transcriptional fusions span the region from the TSS (+1) and up to the same upstream region from the translational fusions and result in constructs of 315 bp (*cobQ*), 220 bp (*cobU*), 247 bp (*cobE*), 282 bp (*cobF*), and 269 bp (*cobH*), respectively. Short transcriptional fusions span the first 100 bp upstream of and include the TSS. Transcriptional fusions were fused to the standardized RBS1 derived from *S. meliloti sinI*. **(B, C)** Activities of long (B) and short (C) transcriptional promoter probes of *E. adhaerens* LU2703 *cob* promoters with and without the addition of 1 mM GB after 6 h of growth in MOPS-buffered minimal medium in microplates. Data is based on at least 4 biological replicates. Statistical significance was calculated by multiple *t* test (n.s.: not significant, *: $p_{adj} < 0.05$, **: $p_{adj} < 0.001$).

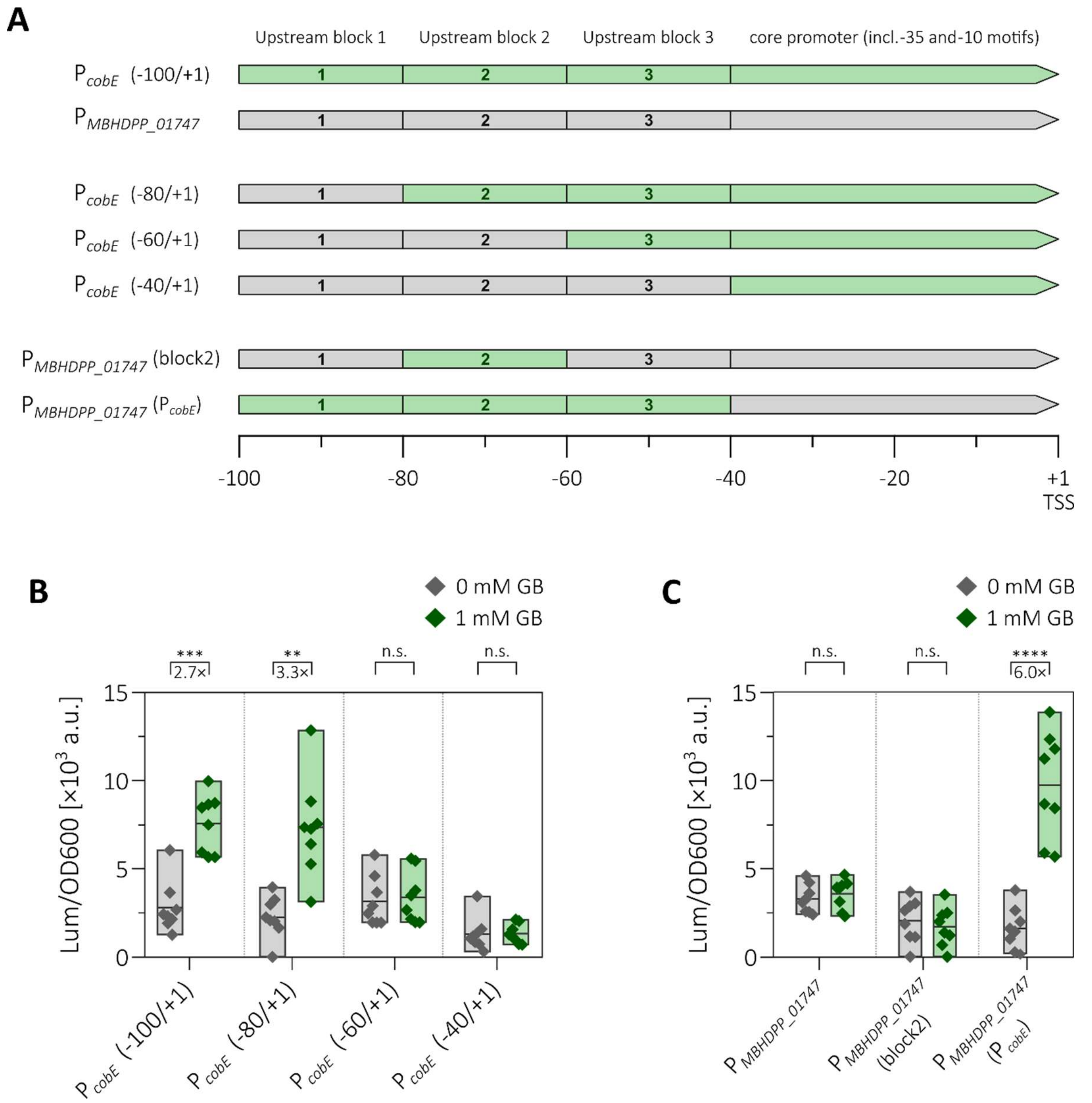


**Fig. S12: Successive replacement of 100 nt *E. adhaerens* LU2703 *cobE* promoter fragments with respective sequences from a promoter not responding to GB ($P_{MBHDPP\_01747}$). (A)** Schematic representation of the design of (hybrid) promoter probes. GB-responsive *cobE* promoter (green) and GB-neutral control promoter $P_{MBHDPP\_01747}$ (grey) were divided into 4 sections: the core promoter up to 40 nt upstream of the TSS which includes -35 and -10 promoter elements, and 3 upstream blocks spanning 20 nt each. Respective replacement variants were designed by successive replacements of upstream blocks from the GB-responsive *cobE* promoter with sequences from the control promoter. Additionally, two promoters were designed with the GB-neutral core promoter and either the replacement of upstream block 2 or the entire upstream region. **(B, C)** Promoter-probe activities of hybrid *cobE* promoters (B) and hybrid *MBHDPP_01747* promoters (C) in *E. adhaerens* LU2703 6 h after addition of 0 mM (grey) or 1 mM GB (green) in MOPS-buffered minimal medium. Cultures were grown in microplates and measured on a Tecan Infinite M200 Pro reader. Data shown is based on 4 biological replicates per strain measured twice on different days. Statistical significance was calculated by multiple *t* test (n.s.: not significant, **: $p_{adj} < 0.001$, ***: $p_{adj} < 0.000\ 1$).

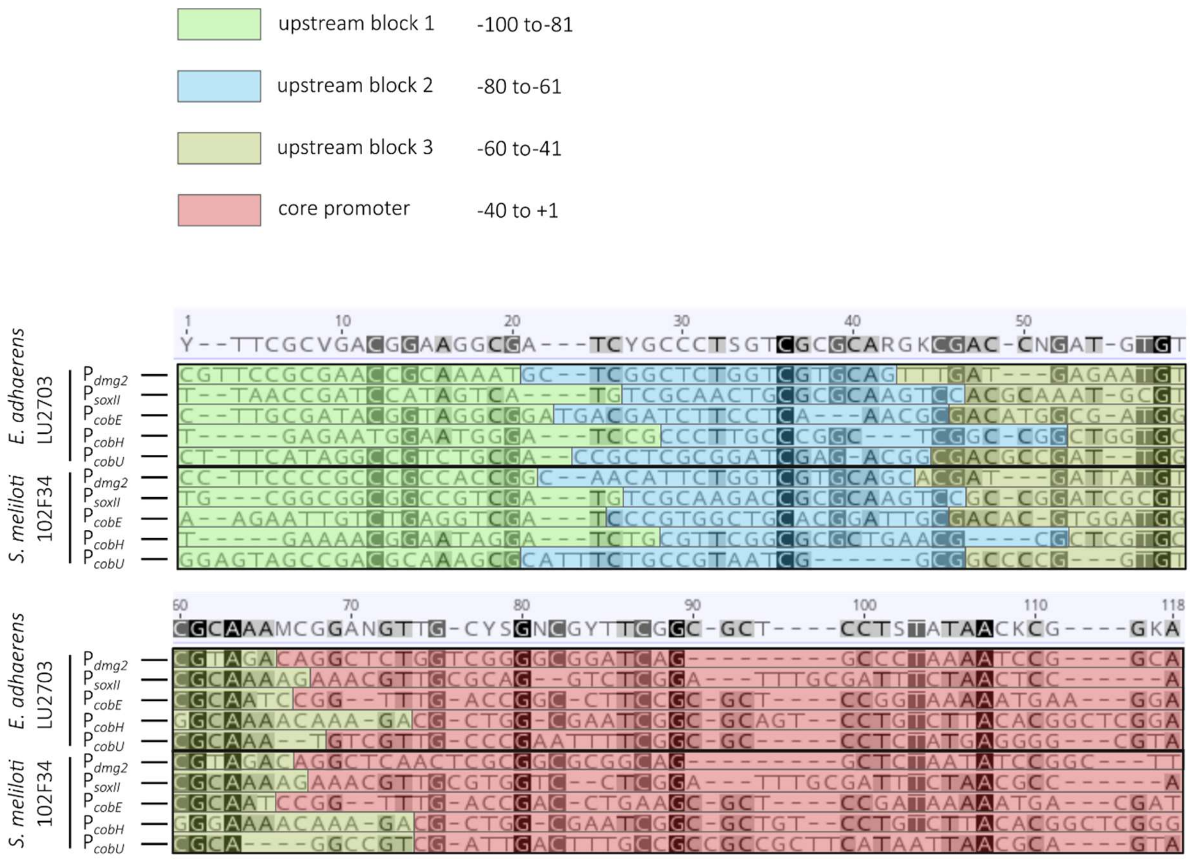


**Fig. S13: Alignment of 100 bp promoter fragments that are transcriptionally regulated by GB in both *E. adhaerens* and *S. meliloti*.** Colored blocks mark 20 bp upstream regions (green, block 1, -100 to -81; blue, block 2, -80 to -61; yellow, block 3, -60 to -41) or the core promoter containing sigma factor recognition motifs (red, -40 to +1).

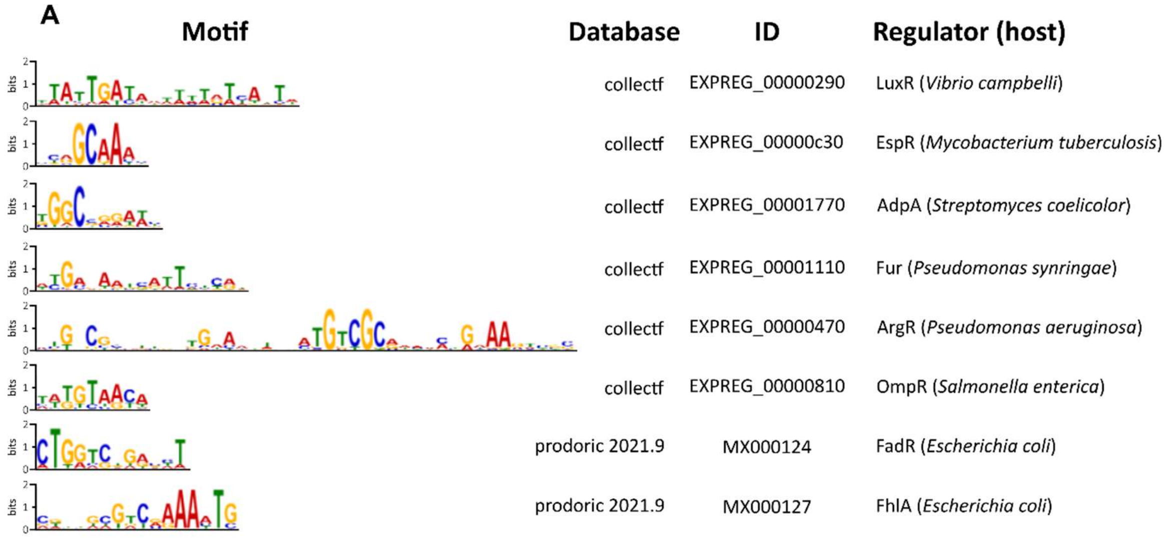


**B**

| SEA hit | Regulated process | % identity of best BLAST hit |
|---|---|---|
| LuxR | Quorum sensing | 46.2% |
| EspR | Pathogenicity | 45.0% |
| AdpA | Secondary metabolism and morphological differentiation | 50.5% |
| Fur | Iron uptake | 43.8% |
| ArgR | Arginine catabolism | 37.0% |
| OmpR | Response to osmotic and acidic stress | 44.5% |
| FadR | Fatty acid metabolism | 29.1% |
| FhlA | Hydrogen production | 41.1% |

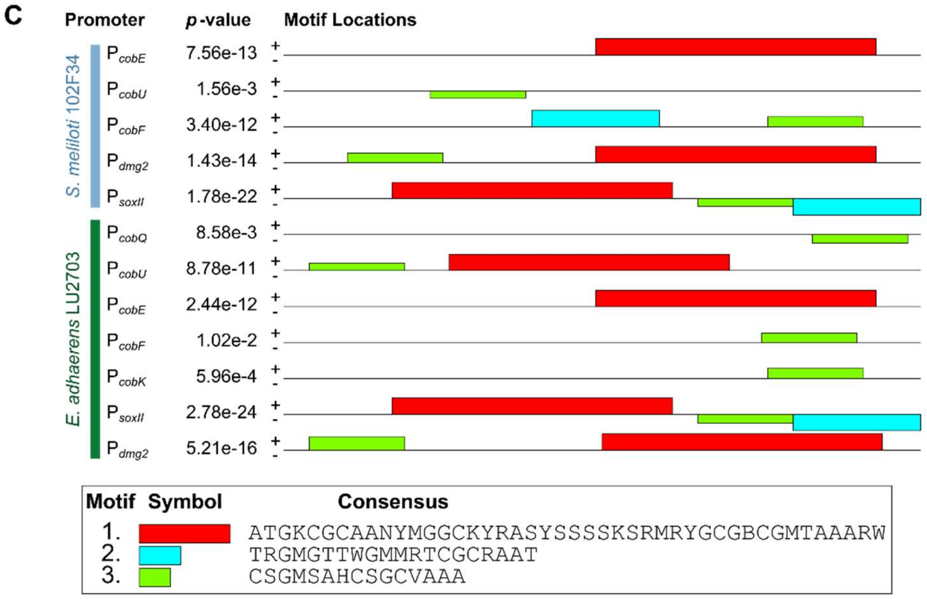


**Fig. S14: Results of motif discovery search in GB regulated promoters of *S. meliloti* 102F34 and *E. adhaerens* LU2703.** Motif search was performed with the MEME software suite and based on 14 GB-regulated promoter fragments using the 100 bp upstream of the respective TSS. Specifically, the promoters of *cobE*, *cobU*, *cobH*, *cobF*, *dmg2*, and *soxII* of *S. meliloti,* and *cobQ*, *cobU*, *cobE*, *cobF*, *cobH*, *cobK*, *soxII*, and *dmg2* of *E. adhaerens* were searched. **(A)** Simple enrichment analysis (SEA) against three bacterial databases (collectf, 84 motifs; prodoric 2021.9, 333 motifs; regtransbase.meme, 141 motifs) revealed potential binding motifs of bacterial transcription factors in the promoters. No motifs were retrieved from the regtransbase.meme database. **(B)** Functional overview of the transcriptional regulators identified in (A). A BLASPp search against the *E. adhaerens* LU2703 proteome revealed proteins with 29.1 % to 50.5 % identity. **(C)** MEME motif discovery search using default settings. No consistent motifs were discovered in the promoter sequences of *cobH* from *S. meliloti* or *E. adhaerens*.

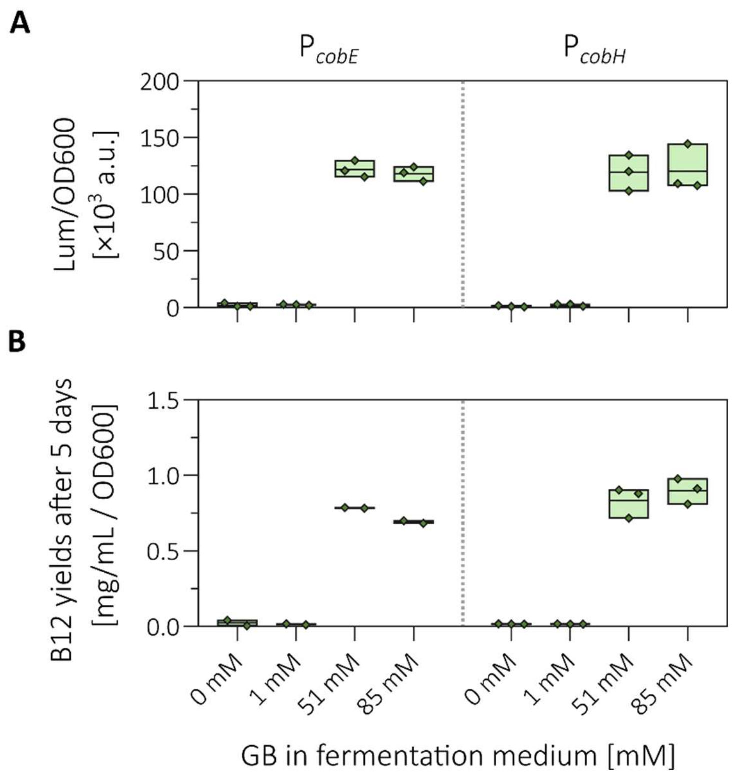


**Fig. S15: Promoter activities and production of vitamin $B_{12}$ by *E. adhaerens* LU2703 at the low and high GB concentrations.** *E. adhaerens* LU2703 was equipped with reporter plasmids allowing either the monitoring of the *cobE* or the *cobH* promoter (same as in Fig. 5). Cultures were grown in MOPS-buffered fermentation medium for 5 days under microoxic conditions in narrow-necked shaking flasks. **(A)** Cultures were diluted 1:100 and OD600 and luminescence was measured using a Tecan Infinite M Plex reader. Promoter activities of *cobE* and *cobH* promoters show no induction at 1 mM after fermentation. **(B)** Quantification of vitamin $B_{12}$ yield via HPLC showed that 1 mM was also insufficient to initiate effective vitamin $B_{12}$ synthesis.

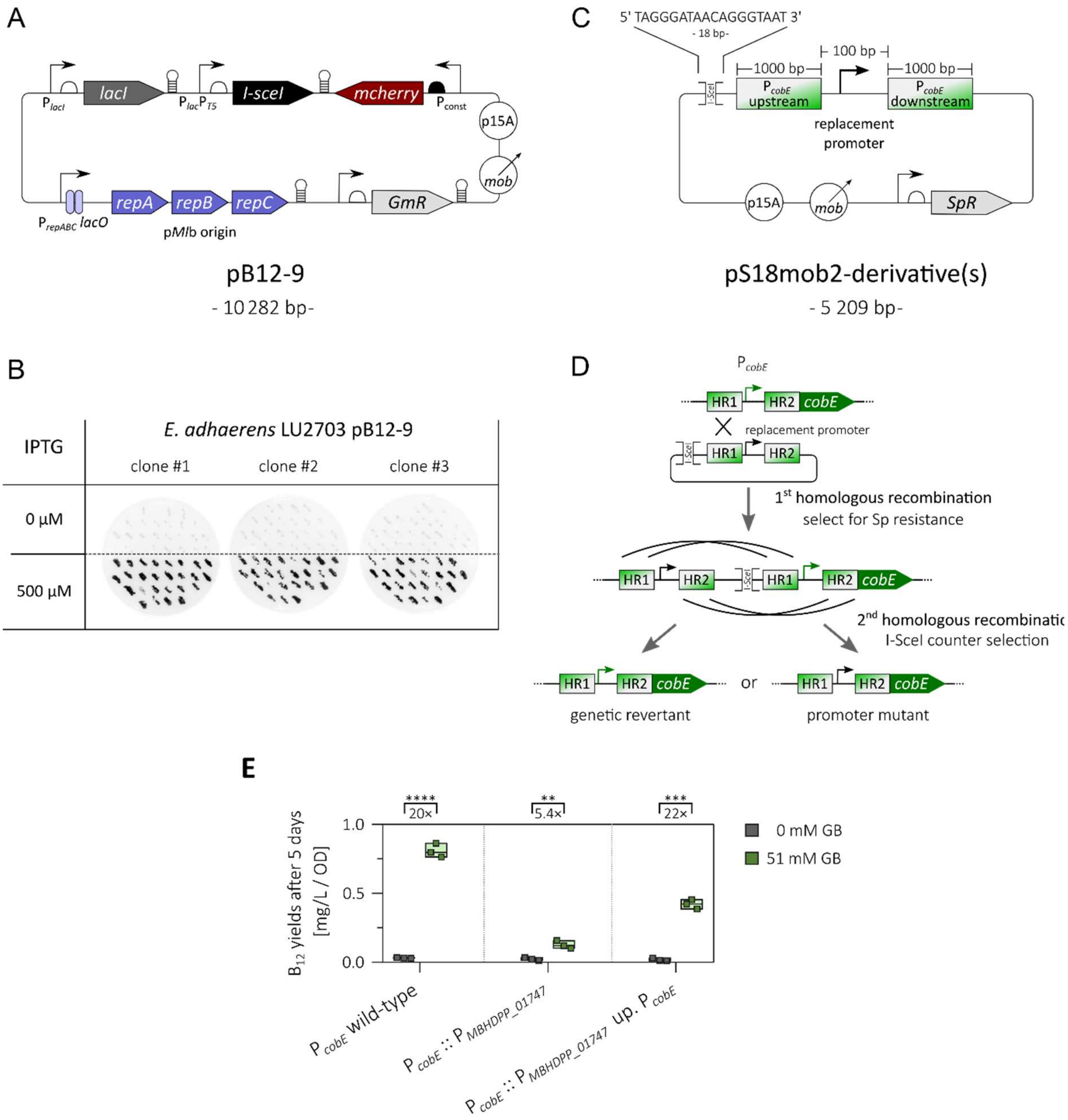


**Fig. S16: An IPTG-dependent counter-selection plasmid expressing the homing endonuclease I-SceI. (A)** Plasmid map of pB12-9, a mobilizable IPTG-dependent *repABC*-type vector based on the p*Ml*b origin of replication. pB12-9 carries the homing endonuclease coding gene *I-sceI* under the control of the IPTG-inducible $P_{lacT5}$ promoter and a constitutively expressed *mcherry* cassette for detection of plasmid presence. It also carries a p15A origin for propagation in *E. coli*. **(B)** Stability of pB12-9 in *E. adhaerens* grown with or without IPTG. pB12-9 was conjugationally transferred to *E. adhaerens* using the standard procedure. Three clones from the same conjugational event were picked and cultured in TY Str600 without gentamycin containing either 0 or 500 µM IPTG for 2 d at 30 °C, 200 rpm, until cultures reached stationary phase. Then, each culture was plated in dilutions 100 to 10-3 on TY Str600 plates maintaining previous IPTG concentrations. From these plates, $n = 25$ colonies were picked and re-streaked on plates containing no IPTG and imaged using an Amersham ImageQuant 800 detecting fluorescence excited at 525 nm. Fluorescence signal indicated the presence of mCherry in the clones. While no clone grown without IPTG showed any fluorescence above background, all colonies grown with IPTG were fluorescent. This demonstrates that pB12-9 can be quickly and reliably cured from *E. adhaerens* via growth in the absence of IPTG. **(C)** Plasmid map of a pS18mob2 derived suicide plasmid that carries a promoter in between two homology regions (HR) upstream and downstream of the *cobE* promoter as well as a I-SceI recognition site. **(D)** Genetic workflow of double homologous recombination in *E. adhaerens*. **(E)** Production of vitamin $B_{12}$ of *E. adhaerens* LU2703 mutant strains with either wild-typical or chromosomally mutated *cobE* promoters after 5 days of microoxic growth in MOPS-buffered fermentation medium. Data shown represents three cultivations of the wild type strain and three independent biological replicates of each mutant strain. Statistical significance was calculated by multiple *t* test (n.s.: not significant, **: $p_{adj} < 0.001$, ***: $p_{adj} < 0.000\,1$, ****: $p_{adj} < 0.000\,01$).
